# Native community resistance modulates the spread of non-native species along Mediterranean mountain roads under global change

**DOI:** 10.1101/2025.10.31.685799

**Authors:** Greta La Bella, Lucia Antonietta Santoianni, Flavio Marzialetti, Andrea De Toma, Claudia Zitarelli, Luigi Cao Pinna, Angela Stanisci, Alicia T:R: Acosta, Fabrizio Bartolucci, Fabio Conti, Marta Carboni

**Affiliations:** Department of Science, Roma Tre University, Rome, Italy; Department of Biosciences and Territory, EnviXLab, University of Molise, Termoli, Italy; Department of Agricultural Sciences, University of Sassari; National Biodiversity Future Center (NBFC), Piazza Marina 61, 90133 Palermo, Italy; School of Mathematics and Statistics, University of Glasgow, Glasgow, United Kingdom; Apennine Floristic Research Center, University of Camerino—Gran Sasso Laga National Park, San Colombo, 67021 Barisciano, Italy

**Keywords:** biological invasion, Central Apennines, climate change, functional traits, land-use change, MIREN, mountains, native community resistance, non-native species, roads

## Abstract

**Aim:** Climate warming, human modification, and global connectivity are eroding the abiotic barriers (i.e. climatic filtering and low propagule pressure) that have traditionally limited the spread of non-native species in mountain ecosystems. Meanwhile, these global changes are reshaping native plant communities, potentially weakening their biotic resistance to invasion. Yet, the role of native community resistance in modulating the upslope spread of non-native species remains overlooked, potentially masking indirect effects of global change in mountain ecosystems.

**Location:** Central Apennines Mountain range, Italy

**Methods:** Following the Mountain Invasion Research Network (MIREN) protocol, we surveyed vegetation close and far from roads at 60 sites along three mountain roads in the Central Apennines. Using current and historical climatic data and aerial images from the 1950s to today, we quantified climate and land-use changes. Dominant plant traits and functional diversity were used to capture the biotic resistance of native communities. We then applied structural equation modelling to investigate how native biotic resistance regulates the effect of global change drivers, i.e. road disturbance and changes in land-use and climate, on the occurrence and cover of non-native species.

**Results:** Variance partitioning suggests that, beside climatic filtering, the biotic resistance of native communities is among the strongest drivers of non-native species occurrence and abundance. Although climate and land-use changes have had little direct influence on non-natives, land-use changes occurred over the past 70 years indirectly influenced invasions by altering native community resistance. Agricultural abandonment at low- to middle elevations, favoured short-statured native communities, offering low resistance. In contrast, forest expansion strengthened resistance through dominance of conservative native species.

**Main conclusion:** Native community resistance, alongside climate and disturbance, is a key determinant of invasion patterns along elevational gradients in Mediterranean mountains. Despite global changes facilitate non-native plant upward shifts, native communities can modulate this spread and may still maintain a certain resistance to invasion.

## INTRODUCTION

Mountains are characterized by unique biodiversity and provide crucial ecosystem services, including provisioning of food and water, carbon storage as well as space for recreational activities (Mengist et al., 2020; Schirpke et al., 2021). However, these environments are particularly sensitive to ongoing global changes, such as warming temperatures, shifting precipitation patterns, and the modification of natural habitats by human activities (Alexander et al., 2016; Dainese et al., 2024; Lembrechts et al., 2016; Pauchard et al., 2009). Yet, until recently, mountain ecosystems had been generally considered to be relatively resistant to another big agent of global change, i.e. the invasion of non-native plant species introduced by humans. This resistance is largely attributed to a combination of low propagule pressure of non-native species, a limited pool of non-natives adapted to the extreme climate of high elevations (i.e. climate filtering), little anthropogenic disturbance, and high resistance to invasion of native communities (Alexander et al., 2016; Milbau et al., 2013; Pauchard et al., 2009). However, climate warming and increased human modification have been reshaping native communities (Collins et al., 2022; Dainese et al., 2017; Geppert et al., 2021), thus potentially altering their resistance to invasion, while simultaneously creating more favourable conditions for the spread of non-native species in mountain areas (Cao Pinna et al., 2024; Petitpierre et al., 2016). However, despite its theoretical importance, the role of native community resistance in modulating the effect of global change on plant invasion in mountain environments remains poorly explored (Alexander et al., 2015, 2018). As the annual rate of new non-native plant introductions continues to grow globally (Seebens et al., 2017, 2021), understanding how climate and land-use changes affect the resistance of native communities is crucial for anticipating and mitigating the spread on non-native species in these vulnerable mountain environments.

The ability of native communities to resist the establishment and expansion of new species explains why, under similar abiotic conditions and propagule availability, a non-native species may or may not successfully invade an ecosystem (Ibáñez et al., 2021; Richardson & Pyšek, 2006). As a result, understanding which features make a community resistant to invasion and how global changes could potentially affect these characteristics is crucial for planning effective management strategies to counteract plant invasion (Roy et al., 2024). In this context, functional traits are increasingly being used as proxies for capturing the ability of native communities to withstand invasions in a wide range of ecosystems (Conti et al., 2018; Gallien & Carboni, 2017) including mountains (McDougall et al., 2018). For example, native communities exhibiting traits related to high resource-use efficiency, strong competitive ability, and rapid growth rates have been shown to resist invasion by limiting the resources accessible to the invaders (Emery & Gross, 2007). Similarly, previous studies demonstrated that communities with greater functional diversity are more likely to exploit a wider range of resources (e.g., light, nutrients, and water) and potentially leave fewer opportunities for invaders to establish (Elton, 1958; Funk et al., 2008). Although some progress has been made in understanding the mechanisms that allow native species to limit the establishment of non-natives, how ongoing global changes are influencing this biotic resistance, and in turn invasion success, still remains poorly explored in mountain ecosystems (but see <u>Barros et al., 2025)</u>.

Climate change, land-use modification, and native community resistance are interconnected forces shaping invasion dynamics (Pyšek et al., 2020). As an example, warming temperature can reduce climatic barriers and allow the expansion of non-native species toward higher elevations (Cao Pinna et al., 2024; Dainese et al., 2017; Pauchard et al., 2016). On the other hand, changes in land-use, such as urban or agricultural expansion, can increase propagule pressure and create new opportunities for non-native plant introduction that can profit from the new favourable conditions (Dainese et al., 2017; Pauchard & Alaback, 2004; Rodewald & Arcese, 2016). Still, changes in climate and land-use do not only influence plant invasion but can also reshape native communities. In particular, climate change has been shown to favour the expansion of thermophilus species from the lowlands (Gottfried et al., 2012; Lenoir et al., 2008; Steinbauer et al., 2022) while some changes in land-use, e.g. the conversion of natural habitat to semi-natural ones, will likely facilitate native ruderal species as an effect of increased disturbance (Tasser & Tappeiner, 2002). As a result, these two drivers of global change do not only influence plant invasions directly, but may also weaken the biotic resistance of native communities, thereby potentially opening new niches for non-native species (Lembrechts et al., 2017a; Pauchard et al., 2016; Petitpierre et al., 2016), or increase biotic resistance instead (Bjorkman et al., 2018). Despite the large body of research that has investigated the effects of climate warming and land-use change on plant invasion in mountain ecosystems (Dainese et al., 2017; Geppert et al., 2021; Pauchard et al., 2009, 2016; Pauchard & Alaback, 2004), the interplay between these drivers of global change and native community resistance and, in turn, their combined effect on plant invasions have been rarely jointly analysed in mountain ecosystems (Alexander et al., 2016)

Mountain roads serve as key corridors for the dispersal of non-native plants along elevation gradients, acting as reservoirs of propagules from nearby lowland and warmer regions (Lembrechts et al., 2017b; McDougall et al., 2018; Pauchard & Alaback, 2004). Additionally, roadsides, often subject to active management practices such as vegetation clearing, provide an opportunity to compare the invasion resistance of disturbed roadside vegetation with undisturbed native plant communities further from the road. As such, mountain roads represent an excellent model system for studying the distribution and abundance of non-native species and their potential for invading high-elevation habitats in the current era of global change. In this study, we aimed to disentangle the complex interactions influencing plant invasion along elevation gradients in Mediterranean mountain ecosystems, focusing on the role played by native community resistance in modulating the effects of global changes on plant invasion. Specifically, (i) we first examined which among road disturbance, climate change, land-use change, and native community resistance is the most influential direct driver of non-native species occurrence and abundance; (ii) then, we explored whether global change drivers indirectly influence plant invasion by altering the resistance of native plant communities.

## MATERIAL AND METHODS

### Study area and sampling design

The study was conducted in the Central Apennines Mountain range of central Italy (Figure S1). The area is characterized by Mediterranean and temperate climate, with a mean annual temperature ranging from 13 °C at the low elevation sites to 4 °C at high altitude sites, and mean annual precipitations varying from 610 to 840 mm along the elevational gradient (Cutini et al., 2021; Pesaresi et al., 2017).

In spring-summer 2022, vegetation was sampled following the MIREN road survey protocol (see Haider et al., 2022 for details about the protocol), which consists in a stratified sampling along mountain roads, so as to cover the main elevational gradient within the mountainous region. Specifically, the sampling was carried out along the road of three mountain sites, namely Mount Terminillo (coordinates at the mountain peak N 42.473958 E 13.066039; elevational range between 420-1915 m a.s.l; Lazio region), mountain chain of Gran Sasso (N 42.442080 E 13.559249; 595-2125 m a.s.l; Abruzzo region), and Mt. Maiella (N 42149692 E 14.115330; 485-2065 m a.s.l; Abruzzo region). These three roads were similarly paved. i.e. asphalt-covered, including at the highest elevations, so road type did not vary across sites. Along each road, we established 20 sites evenly stratified by elevation at approximately 80 m elevation intervals. In each site, we sampled vegetation in two 2 m × 50 m plots: one plot located close and parallel to the roads (hereafter roadside plot), where vegetation is subject to high anthropogenic disturbance; and a second plot perpendicular to the road and distant 50 m from the roadside (hereafter interior plot), representing a vegetation type exposed to lower disturbance levels (Figure S1). In two sites, due to excessive slope, placing the internal plots was not possible. As a result, in total the dataset included 118 plots, 60 roadside plots and 58 interior plots. In each plot, we recorded the presence and relative cover in percentage of all vascular plants. Elevation was measured in the field with a multi-GNSS GPS. Plant species nomenclature followed the Word Flora Online (WFO, 2024). Species were classified into natives and non-natives according to Bartolucci et al. (2024) and Galasso et al. (2018, 2024)

### Plant traits

We used plant traits to represent the competitive ability and functional diversity of the native community in order to capture potential biotic resistance and the availability of open niches for invasion (Gallien & Carboni, 2017). We gathered trait data from the plant trait platform TRY (Kattge et al., 2020). To characterise functional differences between species, we examined four morphological traits responsible of the first two axes of variation in the global spectrum of plant form and function (Díaz et al., 2016). Specifically, we considered two traits associated with plant size, namely plant height (H; m), and seed mass (SM; mg), and two traits related to the leaf economic spectrum, namely specific leaf area (SLA; mm2 mg−1) and leaf nitrogen content (LN, mg g^−1^). These traits reflect the diversity of plant strategies in resource acquisition and competitive ability. Plant size captures the ability to intercept light and dominate the canopy (Violle et al., 2009) while leaf traits are associated with resource use efficiency and relative growth rate (Díaz et al., 2016; Wright et al., 2004). Trait values obtained by TRY were averaged across individuals by species. To deal with missing trait data, we excluded plots where species trait information did not fulfil a threshold of 80% of the total plot cover (Pakeman & Quested, 2007).

### Climate

To reconstruct current climate and climate change occurred over the last 70 years, for the central point between the roadside and interior plot of each site, we gathered monthly precipitation, minimum and maximum temperature data at 1 km resolution for the period between 1931 and 2018 from the CHELSAcruts database (Karger & Zimmermann, 2018). We averaged these data to calculate the mean annual precipitation (MAP), mean annual temperature (MAT), winter precipitation, and summer maximum temperature. These four climatic variables were selected because they capture the main temperature and precipitation gradients relevant to mountain ecosystems and have been shown to be strong predictors of species and ecosystem responses to climate (Barros et al., 2025). These variables were calculated for three different climatic periods, each covering a 30-year interval to account for inter-annual climatic variability (World Meteorological Organization, 2017): T1, running from 1931 to 1960; T2, from 1961 to 1990; and T3, from 1991 to 2020, representing the current climatic period. Although the CHELSAcruts dataset did not include monthly data for the years 2022 and 2020, the climate normal values for the T3 period can be considered reliable because they account for at least 80% of the 30-year interval (World Meteorological Organization, 2017). To increase spatial accuracy, we downscaled the CHELSAcruts climate variables to 30 m resolution using the R package KrigR (Zurell et al., 2021), incorporating digital surface model (DSM at 30 m resolution), slope, and aspect as covariates.

### Land-use

To assess current land-use as well as land-use changes, we visually inspected aerial ortho-photographs with 1 m resolution to produce three land-cover maps of the study area for the years 1954 (T1), 1988-1989 (T2), and 2022 (T3). Land cover was manually photo-interpreted in QGIS environment (QGIS 3.34.11; https://www.qgis.org) within a circular buffer of 250 m of radius around the central point of each site (i.e. 50 m far from the roadside, adjacent to the interior plot). Following the CORINE Land Cover classification, we identified 6 land-cover classes (see Table S1 for a full description): artificial surfaces (ART), roads (ROA), agricultural areas (AGR), afforestations (AFF), forests (FOR), and grasslands (GRA). We thus calculated the percentage landscape (PLand) of each class within each buffer for the three periods, using the ‘lsm_c_pland’ R function (*landscapemetric*; Hesselbarth et al., 2018).

We validated the land cover map relative to the year 2022 (T3) using 200 ground control points collected by field campaigns from 2022 and 2024 and evenly distributed across land-use cover types and the study area. To assess the accuracy of visual inspection, confusion matrices were constructed, and the following performance metrics were calculated: overall accuracy and Cohen’s Kappa statistics (Congalton & Green, 2019). The overall accuracy was 98 % and Cohen’s K = 0.98, indicating a good land cover map accuracy. Since the land cover maps of the years 1954 and 1989 were photo-interpreted using backward editing technique on the basis of the most recent map (2022), the accuracy of older land cover maps can also be considered satisfactory (Malavasi et al., 2013).

### Deriving drivers of invasion

#### Native community resistance

To investigate native community resistance, we used the relative cover of native species (thus excluding non-natives) to calculate functional diversity (FD) and trait community weighted means (CWMs), which quantify respectively, the availability of open niches to invasion and the dominant functional strategy adopted by native species to outcompete non-natives.

For calculating FD, we first computed a multi-trait dissimilarity matrix between all native species using all collected traits (‘gawdis’ R function; de Bello et al., 2021). Then, we employed the functional distance matrix to calculate Rao quadratic entropy using the ‘melodic’ R function (de Bello et al., 2016). We separately calculated the native CWM of plant height, seed mass, SLA, and LN using the **‘**functcomp’ function (R package *FD*; (Laliberté et al., 2014). We then conducted a principal component analysis (PCA, R package *vegan*; Oksanen et al., 2021) using all the CWM values to synthetise dominant native plant form and strategies within each community. The first two PCA axes (Fig. 1d) were used as proxies of native community resistance. The first PCA axis (41 % of the variance) distributes the native communities along a conservative-acquisitive strategy continuum, according to the “slow-fast” leaf economic spectrum (Reich, 2014; Wright et al., 2004), ranging from conservative species characterized by low SLA and LN (negative values) to acquisitive species with high SLA and LN (positive values). The second PCA axis (35 % of the variance) separates native communities according to plant size, from communities dominated by short-statured species with small seeds (negative values) to those dominated by tall species with large seeds (positive values) (Díaz et al., 2016). Here we assume that 1) communities dominated by taller species with larger seeds might be more competitive for light and at the seedling stage and thus have greater resistance to non-native species establishment; 2) similarly communities with high SLA and LN should harbour fast growing species that are potentially more competitive and could also be more resistant to invasion, 3) communities with higher FD should offer less available open niches for non-native species and thus also be more resistant.

**Figure 1.**
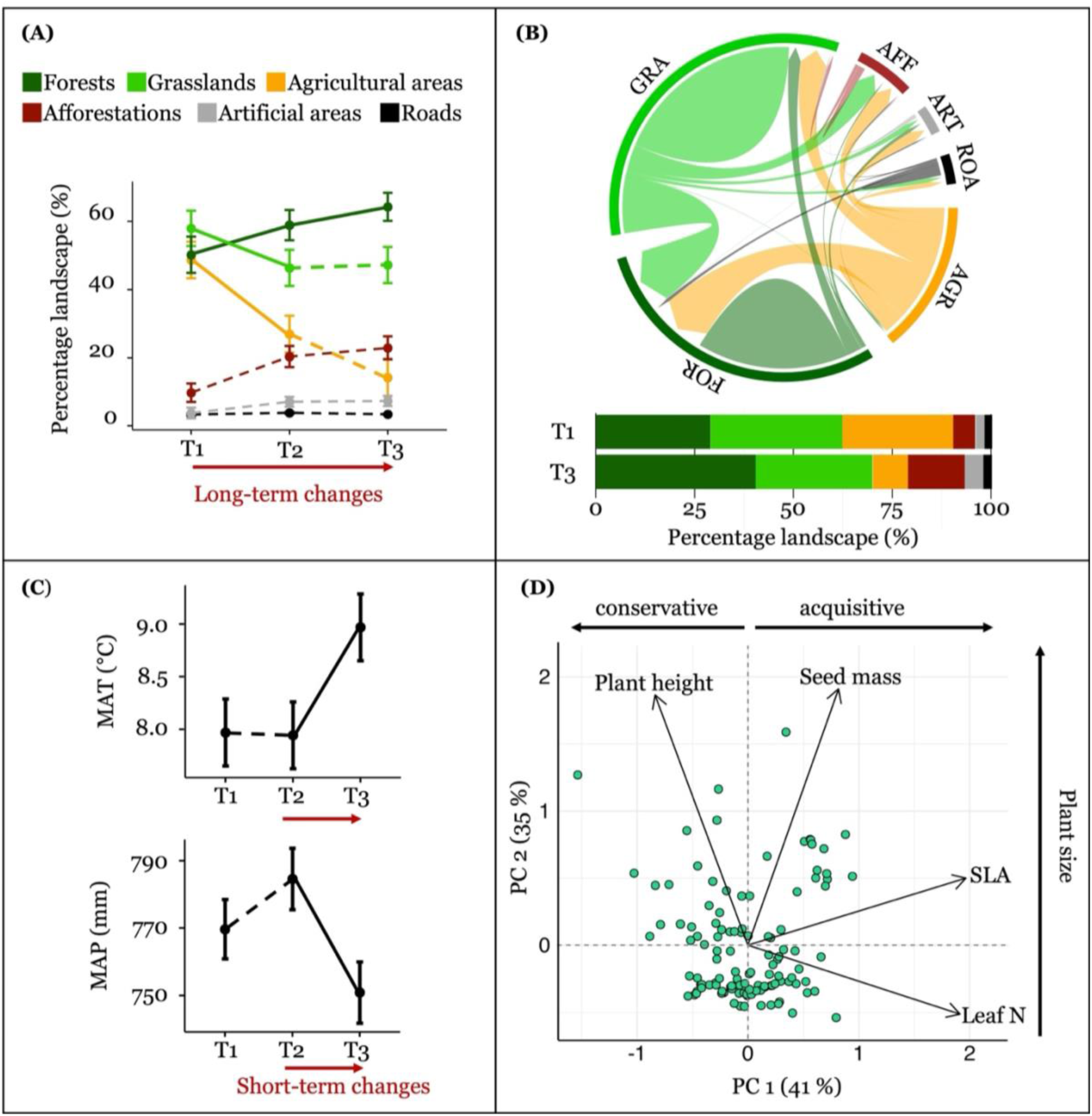
Synthesis of land-use change (A and B), climate change (C), and the native community functional structure (D). (**A**) Changes in the percentage landscape covered by forests, grasslands, agricultural areas, afforestations, artificial areas, and roads occurred over the last 70 years, between 1954 (T1), 1988-1989 (T2), and 2022 (T3). Continuous lines refer to significant changes while dashed lines to non-significant ones (**B**) Cord diagram and bar plot summarizing the changes in percentage of each land cover types between 1954 (T1) and 2022 (T3). In the chord diagram, the arrows represent the direction of change, while the width of the arrow represents the extent (%) of the transition. (**C**) Changes in mean annual temperature (MAT, °C) and mean annual precipitation (MAP, mm) occurred between three 30-year climatic intervals: T1, running from 1931 to 1960; T2, from 1961 to 1990; and T3, from 1991 to 2020. Continuous lines refer to significant changes while dashed lines to non-significant ones (**D**) Principal component analysis (PCA) of the community weighted means (CWMs) of specific plant height, seed mass, leaf area (SLA), and leaf nitrogen content. Each dot refers to a native plant community. The percentage of the total variance explained by each component is given in brackets. The PC1 ordinates the communities along a gradient of conservative-acquisitive dominant strategy while the PC2 separates communities according to plant size.

#### Climate and land-use changes

We first assessed if climate and land use changes actually occurred over the past 70 years in the study area, in order to check if these could be tested as candidate drivers of invasion patterns. Short- and long-term changes in MAT, MAP, and the cover of each land-use classes were calculated, respectively, as the differences between T3 and T2 (Δ_T3-T2_, short-term changes occurred over the last 30-40 years), and the delta between T3 and T1 (Δ_T3-T1_, long-term changes occurred over the last 60-70 years). We thus obtained indices of change with negative and positive values indicating, respectively, a decrease or increase, while values equal to 0 indicate no changes. We then applied mixed-effect models (“lmer” function, lme4 package; Bates et al., 2014) to test if and how climate and land use changed over time. For climate changes, we used MAT and MAP as response variables and “Time” as predictive categorical variable consisting of three levels, i.e. T1, T2, and T3. To test for land-use changes, we used cover as response variable with “Time”, “Land-use cover type”, and their interaction as predictive variables. “Land-use cover type” is a categorical variable consisting of six levels, i.e. artificial surfaces (ART), roads (ROA), agricultural areas (AGR), afforestations (AFF), forests (FOR), and grasslands (GRA). All these models included the site nested in the mountain as random effects in order to account for the repeated measure over time and the variability between mountains. Model residuals were visually inspected to ensure assumptions of homoscedasticity and normality (Zuur et al., 2009). MAT was log-transformed to meet model assumptions. Finally, Tukey’s post hoc tests were used to assess the significance of temporal trends in climate and land-use. We then only considered significant changes as potential drivers of invasion in further analyses, i.e. short-term changes (Δ_T3-T2_) in MAT and MAP, and long-term changes (Δ_T3-T1_) in the cover of agricultural area, forests, and grasslands (see Figure 1a and b).

To better describe and understand these long-term land-use changes, we built a transition matrix summarizing changes occurred in each cover type between T1 and T3 (Malavasi et al., 2018). To do so, we first converted vectorial layers of each cover type into a raster layer with a grain of 1 m for the three time periods. Then we built the transition matrices. Finally we showed these changes using a Chord-Diagram using the R function ‘chordDiagram’ (package *circlize*; Gu et al., 2024).

### Statistical analysis

We used piecewise structural equation modelling (SEM, *piecewiseSEM* package; Lefcheck, 2016) to investigate direct and indirect drivers of non-native species occurrence and abundance. Based on current knowledge about the drivers of plant invasion along elevation gradients, we outlined two meta-models (Figure S2 and Table S2), one for non-native species occurrence and one for non-native species abundance, which involved all hypothesised causal paths linking road disturbance, native community resistance, current climate and land-use, temporal changes in climate and land-use with non-native species occurrence and abundance. Road disturbance, current climate and land-use, as well as temporal changes in climate and land-use are expected to directly influence non-native species occurrence and abundance but also indirectly, by altering the resistance of the native community to plant invasion (Lembrechts et al., 2017a; Pauchard et al., 2016; Petitpierre et al., 2016), here expressed as native plant size, conservative-acquisitive plant strategy, and functional diversity. Finally, we also predicted a direct influence of native community resistance on non-native species occurrence and abundance.

For testing the factors directly driving the occurrence and abundance of non-native species along Mediterranean mountain roads, we fitted two separate generalized linear mixed-effect models (“glmer” R function, *lme4* package; Bates et al., 2014). Specifically, to model non-native species occurrence we used a logistic regression (binomial family) with presence/absence as response variable. To model non-native species abundance, we restricted the analysis to plots where non-native plants were present. Abundance data were expressed as cover proportion and logit-transformed to meet model assumptions. We then fitted a linear mixed-effects model (“lmer” R function, *lme4* package; Bates et al., 2014). In both models, the explanatory variables were: plot disturbance as a two-level factor (roadside plot and interior plot), elevation, plant size (first PCA axis), conservative-acquisitive plant strategy (second PCA axis), functional diversity of native species, current MAT and MAP (during T3 interval), short-term changes (Δ_T3-T2_) in MAT and MAP, current cover of each land-use cover type (at T3), and long-term changes (Δ_T3-T1_) in the cover of agricultural area, forests, and grasslands. Prior to model fitting, we checked for multicollinearity between predictors using the ‘vifstep’ R function (*usdm* package; (Naimi, 2012) and excluded those with a Variance Inflation Factor (VIF) exceeding 3 (Zuur et al., 2009). The final set of predictive variables included: plot disturbance, plant size, conservative-acquisitive plant strategy, functional diversity, current MAT, current MAP, short-term changes in MAT and MAP, current cover of agricultural areas, artificial areas, roads, and grasslands, long-term changes in the cover of agricultural areas, grasslands, and forests. Note that elevation was excluded because, as expected, it was highly correlated with MAT (*r* = 0.99, *p-value* < 0.001). On the other hand, for testing how global change factors indirectly influence plant invasion via changes in the resistance of native communities, we fitted three separate linear mixed-effect models. Response variables were native plant size, conservative-acquisitive plant strategy, and functional diversity. Final predictors after checking for multicollinearity were: plot disturbance, current MAT, current MAP, short-term changes in MAT and MAP, current cover of agricultural areas, artificial areas, roads, and grasslands, long-term changes in the cover of agricultural areas, grasslands, and forests. Considering all these relationships, each SEM consisted of four sub-models (listed in Table S2), all including mountain identity as random effect in order to account for the variability between mountains. SEM goodness-of-fit was evaluated through Fisher’s *C* statistic with *p*-values over 0.05. From the initial SEM, non-significant paths were sequentially deleted, and the final structure was selected based on Akaike Information Criterion (AIC < 2; (Burnham & Anderson, 2002; Richards, 2005; Shipley, 2013). Model assumptions were visually inspected. We finally computed indirect effects as the product of the coefficients of the paths that form the indirect relationship (Gana & Broc, 2019).

To assess the most important drivers of non-native species occurrence and abundance, we performed a variance partitioning on the final models using the *variancePartition* R package (Hoffman & Schadt, 2016). This approach quantifies the relative importance of each predictor as the percentage of the total variance explained by the model.

All statistical analyses were conducted with R version 4.3.1 (R Core Team, 2023).

## RESULTS

### Non-native species

Across the surveyed plots, we found a total of 811 vascular plant species, ranging from 10 to 97 species per plot (mean = 43.63, SD = 16.81). Among these, 16 were classified as non-natives (Table S3<u>; for more details see Santoianni et al., 2025)</u>. Within invaded plots, the number of non-native species ranged between 1 and 4 (mean = 1.77, SD = 0.9), while the non-native species cover ranged between 0.1 % to 45.5 % (mean = 5.43 %, SD = 9.30).

### Changes in climate and land-use

We found significant changes in both climate and land-use in this area over the past 70 years (Figure 1a-c; Table S4). However, changes in climate were more pronounced over the last 30 years (Figure 1c; Table S5), i.e. between the climatic periods 1961-1990 (T2) and 1991-2020 (T3), while changes in land-use occurred earlier in time (Figure 1a; Table S6), starting from 1954 (T1). We thus considered only short-term changes for climate (Δ_T3-T2_) and long-term changes for land-use (Δ_T3-T1_) in further analyses. Between T2 and T3, MAT increased (t = 0.897 °C, *p-value* < 0.001) while MAP decreased (t = -30.0 mm, *p-value* < 0.001). On the other hand, between 1954 (T1) and 2022 (T3), the study area experienced a decline in the cover of agricultural areas (t = -34 %, *p-value* < 0.001) and grasslands (t = -10 %, *p-value* < 0.048) that have been replaced by forests’ expansion (t = 14 %, *p-value* < 0.017, Figure 1b).

### Most important drivers of invasion

Through variance partitioning we identified the most important drivers influencing non-native species occurrence and abundance (Figure 2). Overall, both models demonstrated good explanatory power (R^2^), explaining 65 % of the variation in both non-native apecies occurrence and abundance. We found that non-native species occurrence was mainly driven by climate, with current MAT and MAP explaining 47 % and 6 % of the total variance, respectively, for a combined 53 %). Road disturbance played a secondary role, explaining only 30 % of the total variance. Native community resistance also contributed, although to a lesser extent, with plant size accounting for 10 % and conservative-acquisitive strategy for 6 %, together explaining 16 % of the total variance. On the other hand, the abundance of non-native species was mainly influenced by native community resistance and climate. Plant size, conservative-acquisitive strategy, and functional diversity explaining 14%, 4%, and 10% of the variance, respectively, for a total of 28 %, while current MAT and MAP explained 12 % and 13 % of variation, for a total of 25 %. Temporal changes in climate and land-use, which accounted for none or very little of the variation in non-native species occurrence (only 1% of changes in agricultural areas), had a clearer effect on abundance. Specifically, short-term changes in MAT accounted for 11 % of the variation, while long-term changes in surrounding land-use overall explained 19 % of variation (12 % due to changes in agricultural areas and 7 % due to changes in forest cover). Finally, road disturbance had a negligible effect, explaining only 2 % of variance in non-native species abundance.

**Figure 2.**
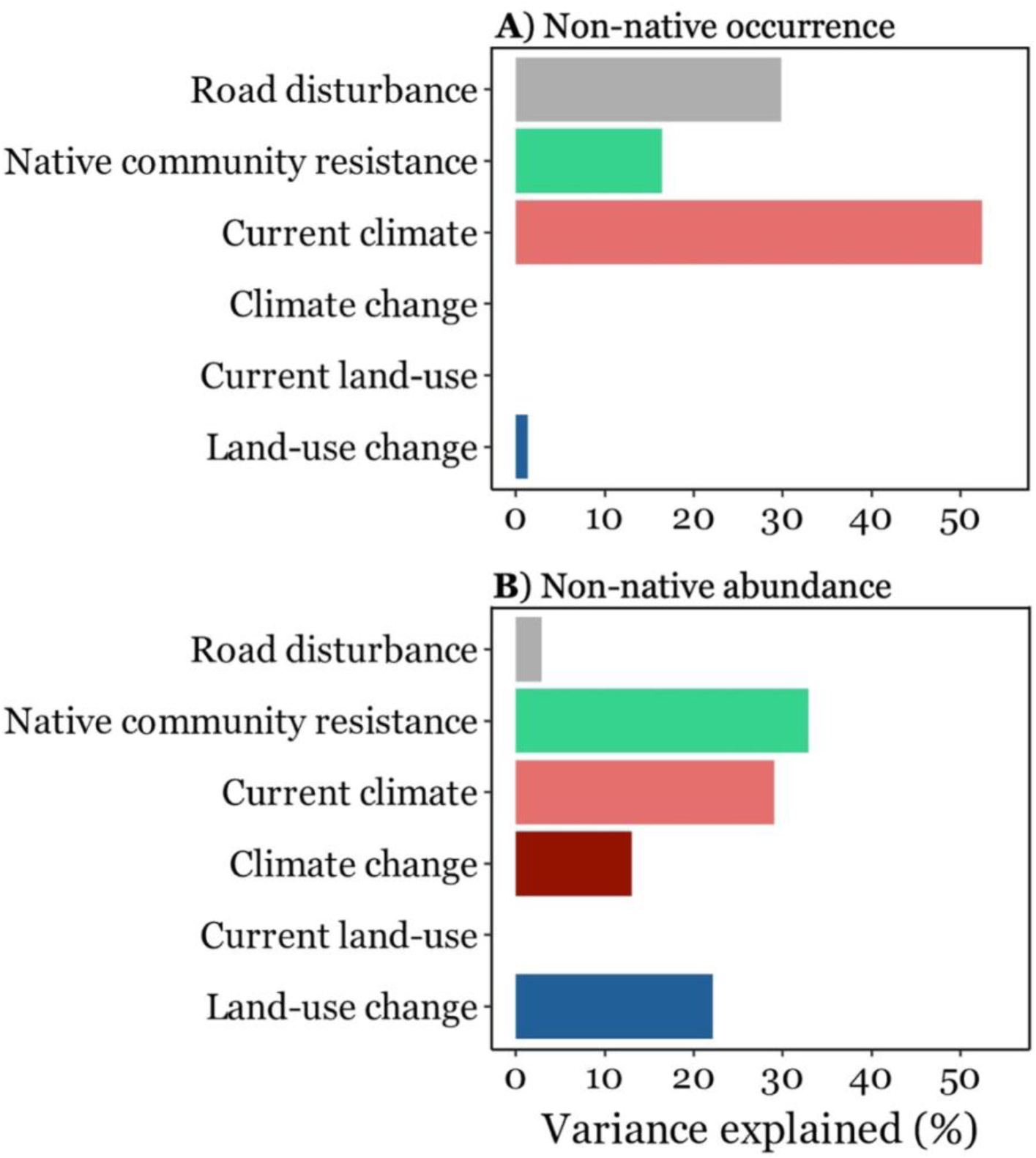
Variance partitioning of the drivers of non-native species occurrence (A) and abundance (B). The relative importance is expressed as % of the total variance explained (model *R*^2^) by each predictor in the models. Native community resistance (in green) refers to the summed effect of dominant plant size, conservative-acquisitive strategy, and functional diversity of the native communities. Current climate (in light red) is the summed effect current mean annual temperature (MAT) and precipitation (MAP), while climate change (in dark red) is the summed effect of short-term changes in MAT and MAP occurred over the last 40 years (Δ_T3-T2_). Current land-use (in light blue) includes the effects of the percentage landscape (PLand) covered by agricultural areas, artificial areas, roads, and grasslands at T3, while land-use change (in dark blue) refers to the summed effects of long-term changes occurred in the surrounding landscape over the last 70 years (Δ_T3-T1_) in the cover of agricultural areas, grasslands, and forests.

### The direct effects of global change and native community resistance on plant invasion

Both SEMs demonstrated a good fit to the data (SEM of non-native species occurrence Fischer’s C = 49.06, df = 50, p-value = 0.511; SEM of non-native species abundance Fischer’s C = 28.71, df = 34, p-value = 0.724; Figure 3) and identified the direct and indirect drivers of non-native species occurrence and abundance. Road disturbance had a positive direct effect on the occurrence of non-native species (std. estimate = 2.72, *p-value* < 0.001; Figure 4a) but no effect on their abundance (std. estimate = 1.13, *p-value* = 0.071; Figure 5a), indicating higher probability of invasion in roadside-disturbed plots compared to interior-undisturbed plots, but similar abundance. Native plant size had a negative effect on non-native species occurrence (std. estimate = -0.790, *p-value* < 0.024; Figure 4b) while native conservative-acquisitive strategy had a positive one (std. estimate = 0.63, *p-value* < 0.048; Figure 4c), denoting higher probability of invasion in native communities dominated by short-statured plant species with an acquisitive resource strategy. We also found higher non-native species occurrence in sites with higher MAT (std. estimate = 1.71; *p-value* < 0.001; Figure 4d), while MAP had no effect (std. estimate = -0.59, *p-value* > 0.05; Figure 4e). On the other hand, the abundance of non-native species was similarly influenced by native community functional structure, with a negative effect of native plant size (std. estimate = -1.28, *p-value* = 0.002; Figure 5b) and a positive effect of native functional diversity (std. estimate = 1.07, *p-value* = 0.020; Figure 5d). Moreover, we found that non-native species were more abundant in sites with lower MAT (std. estimate = -1.15, *p-value* = 0.002; Figure 5e) and MAP (std. estimate = -1.23, *p-value* = 0.003; Figure 5f) as well as sites where MAT increased the most over the last 40 years (std. estimate = 1.12, *p-value* = 0.031; Figure 5g). Finally, non-native species abundance was higher in sites that over the last 70 years experienced in the surrounding landscape a contraction in the cover of agricultural areas (std. estimate = 1.19, *p-value* = 0.016; Figure 5j) and an expansion in forest cover (std. estimate = 0.86, *p-value* = 0.007; Figure 5h).

**Figure 3.**
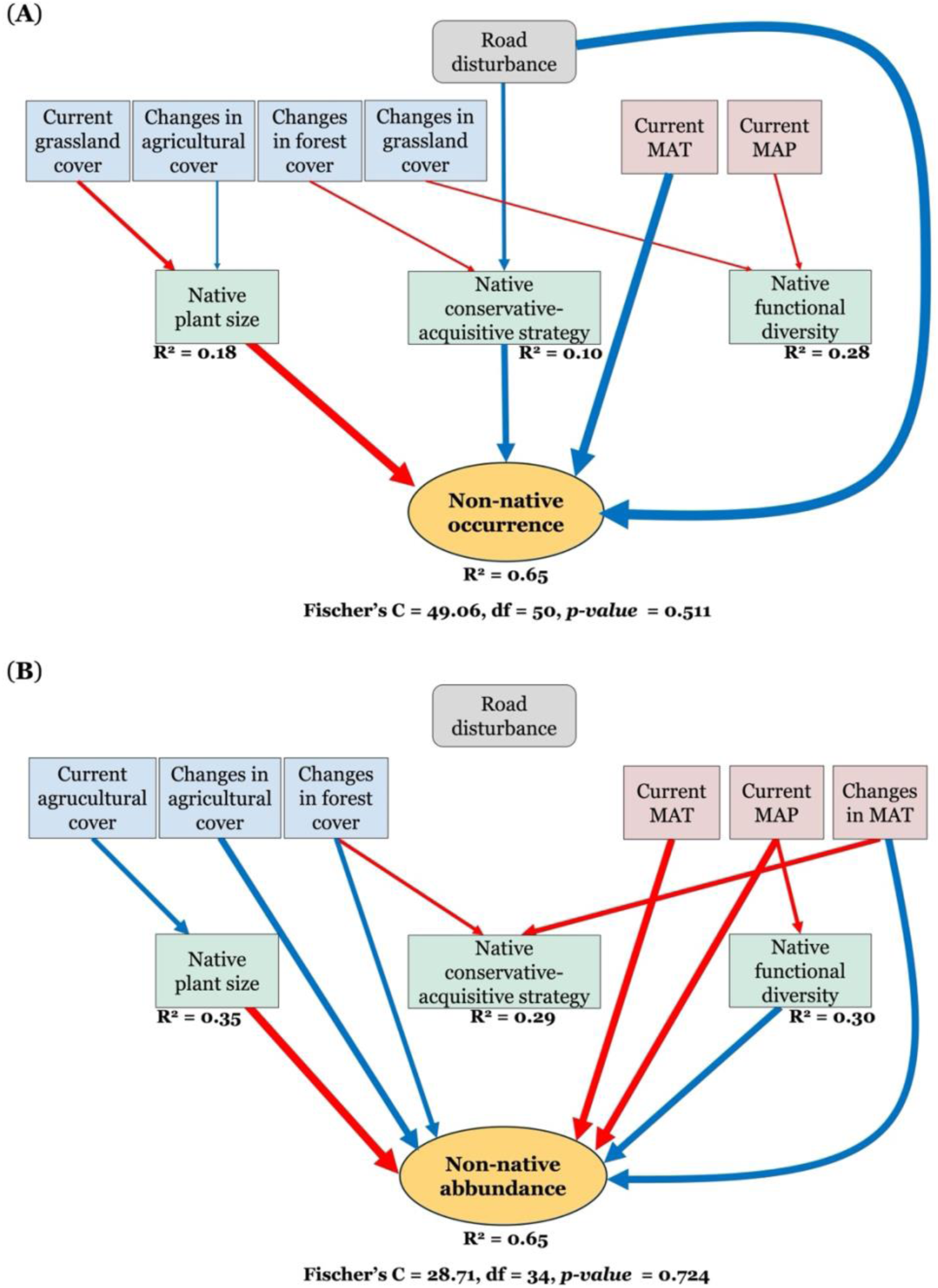
Direct and indirect drivers of plant invasion. Structural equation model (SEM) illustrating the effects of road disturbance, native community resistance, current climate, land-use as well as changes in climate and land-use, non-native species occurrence and abundance and the interplay between these drivers. Red arrows refer to negative effects and blue arrows to positive ones. Arrows’ width is scaled by standardised estimates. R^2^ are reported next to the corresponding response variable.

**Figure 4.**
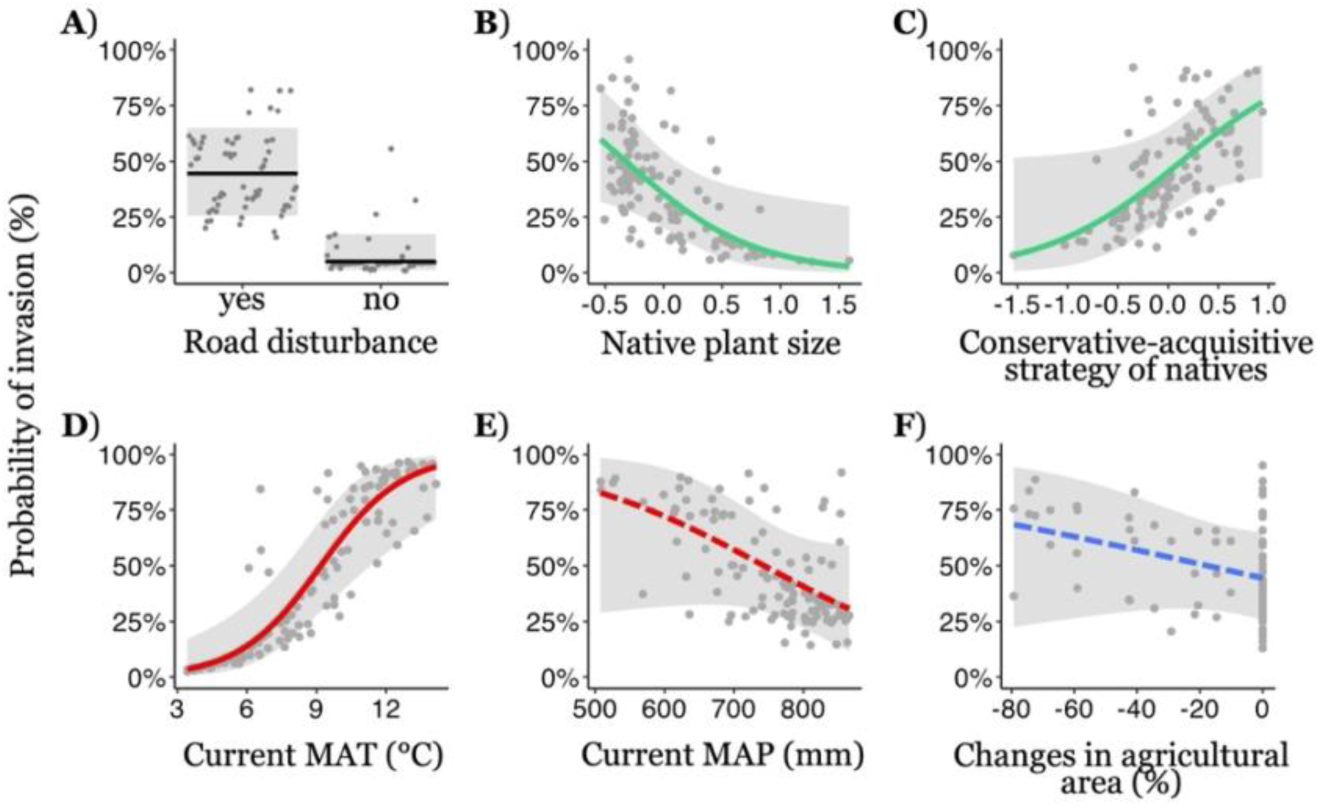
Direct drivers of non-native species occurrence. The effects of road disturbance (A), native community resistance (B and C, in green), current climate (D and E, red), and changes in land-use (F, in blue) on non-native species occurrence, expressed as probability of invasion in percentage. The figure only shows final predictors retained after model selection of the generalized linear mixed-effect model testing the drivers of non-native species occurrence. Continuous regression lines refer to significant effects (*p-value* < 0.05), while dotted lines to marginal and non-significant effects (*p-value* > 0.05). shading areas associated with the lines represent the 95% confidence interval.

**Figure 5.**
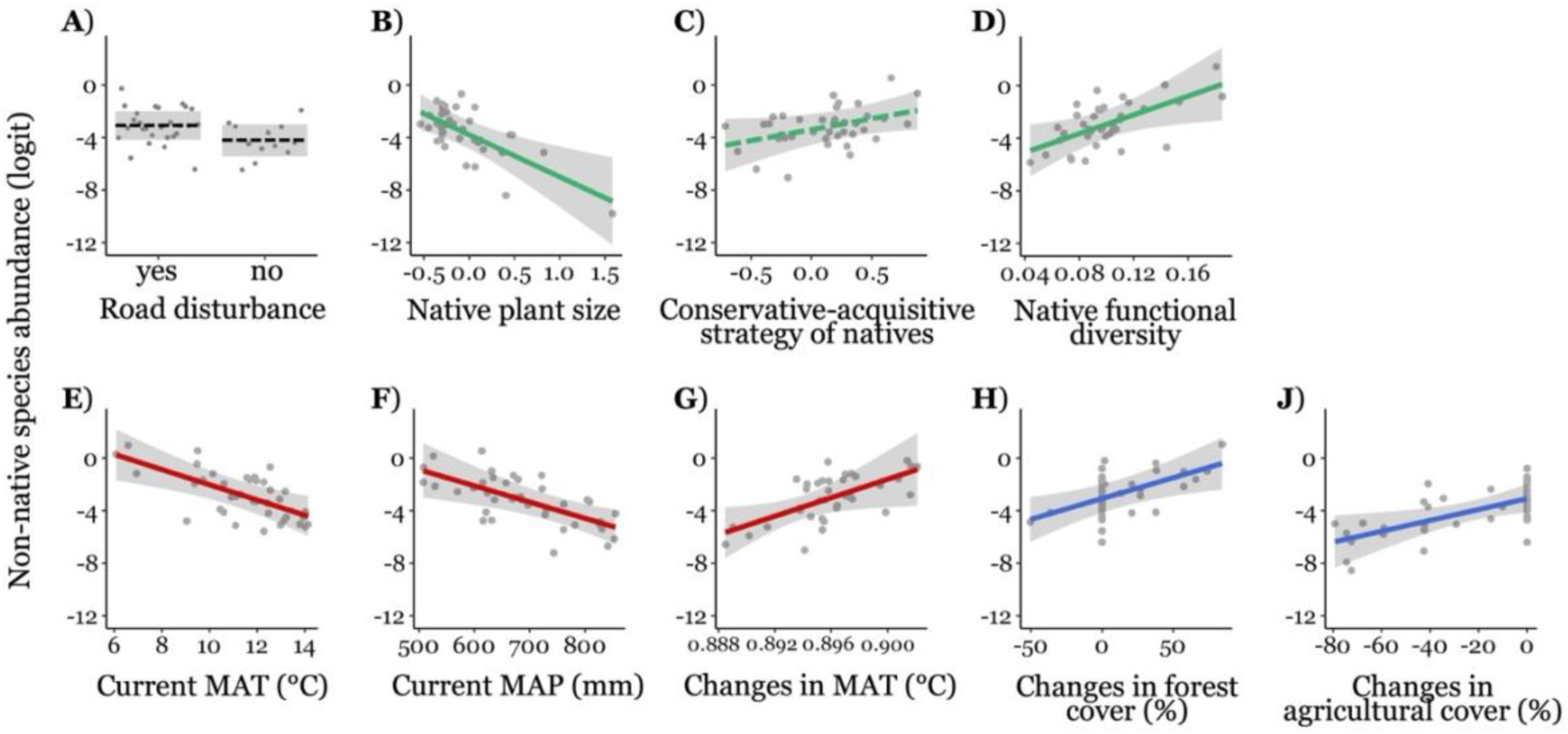
Direct drivers of non-native species abundance. The effects of road disturbance (A), native community resistance (B-D, in green), current climate (E and F, red), and changes in climate (G, in red) and land-use (H and J, in blue) on non-native species abundance (logit). The figure only shows final predictors retained after model selection of the linear mixed-effect model testing the drivers of non-native species abundance. Continuous regression lines refer to significant effects (*p-value* < 0.05), while dotted lines to marginal and non-significant effects (*p-value* > 0.05). shading areas associated with the lines represent the 95% confidence interval.

### The indirect effects of global change on plant invasion modulated by native community resistance

Through the SEM analysis, we showed that global change factors also indirectly affect non-native plants by altering the resistance of native communities to invasion (Figure 3). Global change drivers have indeed led to significant shifts in the native communities. Sites that experienced a decrease in agricultural cover were dominated by short-statured species with small seeds (std. estimate = 0.23, *p-value* = 0.036; Figure 3a), which in turn increase invasion probability (indirect effect = -0.18; Figure 3a). In contrast, forest expansion indirectly reduced invasion probability (indirect effect = -0.15; Figure 3a) by promoting the dominance of conservative species within native communities (std. estimate = -0.24, *p-value* = 0.014; Figure 3a), which are more resistant to invasion. Higher grassland cover in the surrounding landscape indirectly promoted non-native species occurrence (indirect effect = 0.30; Figure 3a), as it was associated with smaller native plant size (std. estimate = -0.38, *p-value* = 0.001; Figure 3a), and thus with lower biotic resistance. Road disturbance favoured the dominance of acquisitive strategies (std. estimate = 0.39, *p-value* = 0.029; Figure 3a), thereby indirectly promoting non-native species occurrence (indirect effect = 1.06; Figure 3a). By contrast, the abundance of non-native species was negatively influenced indirectly by both land-use and climate. Current agricultural cover increased native plant size (std. estimate = 0.47, *p-value* = 0.009; Figure 3b), which in turn decreased non-native species cover (indirect effect = 0.60; Figure 3b). Likewise, current MAP had a negative indirect effect on non-native species abundance (indirect effect = -0.42; Figure 3b) through its negative effect on native functional diversity (std. estimate = -0.39, *p-value* = 0.039; Figure 3b.

## DISCUSSION

Native community resistance plays a central role in regulating the establishment and spread of non-native species across mountain ecosystems, yet its importance is often overlooked in favour of direct global change drivers. While human disturbance, climate and land-use changes are expected to facilitate plant invasion across mountain ecosystems (Alexander et al., 2016; Carboni, Calderon-Sanou, et al., 2018; Lembrechts et al., 2016; Pauchard et al., 2009, 2016), it remains unclear whether their effect are direct or are instead modulated by changes in the biotic resistance of native communities. This study addressed this question by examining how global change drivers, such as human disturbance (here expressed by road disturbance), climate change, and land-use changes, have affected the occurrence and abundance of non-native species in Mediterranean mountain ecosystems, both directly and indirectly through their impact on the functional structure and composition of the native communities. Our results highlight that although climate filtering remains a key direct drivers of plant invasion, the resistance of the native community is a decisive factor that modulates the distribution and abundance of non-native species along elevational gradients and their response to global changes. For example, the abandonment of agricultural lands has resulted in the development of short-statured native plant communities that exert lower biotic resistance to non-native plants establishment; by contrast, the regeneration of forests on former agricultural lands has strengthened native community resistance through the dominance of conservative species. Overall, these findings demonstrate that in Mediterranean mountain ecosystems, native communities continue to provide significant resistance to plant invasion despite ongoing global changes. This highlights the importance of maintaining and restoring native communities to mitigate future invasion risks.

### The influence of global change drivers and biotic resistance on plant invasion

High elevation plant communities are still considered to be less affected by plant invasion thanks to a combination of climate filtering, low propagule pressure, and limited human disturbance (Alexander et al., 2016; Dainese et al., 2024; Lembrechts et al., 2016; Pauchard et al., 2009). In support of this idea, mean annual temperature (MAT) emerged as the strongest driver of the presence of non-native species across the mountains of the Central Apennines, explaining 47 % of variation in non-native species occurrence (Fig. 2). This finding underscores the critical role of temperature as a limiting factor for non-native species establishment at higher elevations, where colder conditions and shorter growing seasons act as strong climatic filters (Hoch & Körner, 2012; Sundqvist et al., 2013). Although most studies did not explicitly test the direct influence of temperature in driving non-native species patterns across mountain ranges, often using elevation as a proxy of temperature variation (Körner, 2007), similar patterns of declining non-native species diversity with decreasing temperature have been consistently observed worldwide (Haider et al., 2018; Pauchard et al., 2009; Seipel et al., 2016). This is because non-native species, often originating from warmer regions (Cao Pinna et al., 2021), face significant challenges in surviving and reproducing in colder, high-altitude environments where metabolic processes and resource availability are constrained (Hoch & Körner, 2012; Sundqvist et al., 2013). Nevertheless, the effect of climate on non-native species abundance was lower than on occurrence. While MAT and MAP together explained about half of the variation in non-native species occurrence, they accounted for only about a quarter of the variation in non-native species cover, slightly less than the variation explained by native community resistance (28 %). This suggests that although climate remains a critical filter for determining whether non-native species can establish at higher elevations, their success in terms of dominance and spread within communities is more strongly regulated by biotic resistance. In other words, climate sets the stage for non-native species entry, but it is the composition and functional structure of native communities that largely determine their abundance once established. This pattern is consistent with theoretical and empirical evidence that competitive interactions and resource-use complementarity constrain the growth of non-native species (Cavieres, 2021; Funk et al., 2008). Our results therefore highlight the need to consider not only abiotic filters but also the biotic context in predicting the outcomes of plant invasions under ongoing global change.

Alongside temperature, human disturbance, quantified with road disturbance, emerged as a critical driver of non-native plant occurrence but not of their abundance. Our findings revealed a greater probability of invasion in disturbed roadside plots compared to undisturbed ones located farther from roads. This pattern aligns with the well-documented role of roads as key vectors for invasion, facilitating both the dispersal of propagules and the creation of suitable conditions for establishment, such as altered soil microclimate and increased light availability (Haider et al., 2018; Lejeune et al., 2024; Lembrechts et al., 2017a; McDougall et al., 2018). However, once established, the cover of non-native species did not differ significantly between roadside and more distant plots. This suggests that while roads strongly influence the entry of non-native species into mountain ecosystems, their subsequent spread and dominance within communities depend more on local biotic interactions and resistance mechanisms than on disturbance per se. In other words, disturbance acts primarily as a gateway for invasion, whereas the degree of invasion success is modulated by the ecological characteristics of the receiving community (Christen & Matlack, 2009; McDougall et al., 2018).

Linked to human disturbance, we also expected an influence of the cover of artificial structures and road networks in the surrounding on the presence and abundance of non-native species. Surprisingly, however, our analysis did not identify any significant effect of these variables. One possible explanation for this result is that, while artificial structures and road networks can serve as significant sources of propagule introduction and dispersal, their impact on invasion success may be limited to immediate surroundings and become more apparent only at finer spatial scales, where proximity to these features directly increases propagule pressure (Fuentes-Lillo et al., 2021). Another potential explanation involves a time lag in the response of non-native species to human modification of the landscape. While propagules may already be present, the establishment and spread of non-native plants can require time to overcome environmental barriers or take advantage of changes in native community composition (Carboni, Guéguen, et al., 2018). Such delays may obscure the current influence of artificial structures and road networks, even though their long-term role in facilitating invasions may still be significant (Essl et al., 2015). Our findings also show that the direct influence of land-use changes on non-native species was minimal. Nevertheless, their impact was exerted indirectly by altering the composition of native communities, thereby reshaping their biotic resistance to invasion (see below).

Biotic resistance emerged as key determinant of plant invasion. Biotic resistance, driven by interactions within native communities, plays a critical role in limiting the establishment and spread of non-native species (Elton, 1958; Richardson & Pyšek, 2006). Our results highlight how the functional traits and composition of native communities shape their ability to resist invasions. Specifically, we found higher biotic resistance (i.e. lower occurrence and abundance of non-native plants) in communities dominated by tall-statured species with large seeds and a conservative strategy in the leaves. In line with our expectations, these results confirm that higher plant size confers greater resistance to invasion. This is because taller species with larger seeds are more competitive in light acquisition and at the seedling stage (Funk et al., 2008; Simpson et al., 2021; Suárez-Vidal et al., 2017; Violle et al., 2009; Westoby et al., 2002). However, we also assumed stronger biotic resistance in communities with acquisitive leaf traits rather than conservatives. Our assumptions rely on the mechanisms of competitive exclusion and limiting similarity (Daehler, 2001; MacArthur & Levins, 1967; Shea & Chesson, 2002) by which native species with functional traits related to rapid turnover and resource acquisition, like high SLA and LN, are highly competitive, leaving few niches available for invaders to establish and persist non-native species (Kraft et al., 2015; Sheppard, 2019). Since also successful non-native plants are usually acquisitive, fast-growing species (Dawson et al., 2012; Heberling & Fridley, 2013; Richardson & Pyšek, 2006; Van Kleunen et al., 2010), one hypothesis is that they tend to avoid native communities with similar traits that potentially use similar resources (Emery & Gross, 2007; Funk et al., 2008). Accordingly, previous research suggested that native communities with acquisitive strategies are more resistant to plant invasion compared to conservative communities (Carboni et al., 2016; Drenovsky & James, 2010; Van Kleunen et al., 2010). While competitive exclusion and limiting similarity are key mechanisms, in line with the preadaptation hypothesis that stresses the role of environmental filtering (Duncan & Williams, 2002), non-native species may preferentially invade acquisitive communities because their traits better align with the environmental conditions and resource dynamics of these systems. Many studies have already demonstrated the strong influence of similarity between native and non-native species on invasion success in a wide range of environmental conditions (Carboni et al., 2013; Fan et al., 2023; Gallien et al., 2015; Le et al., 2024; Park et al., 2020) as well as the higher probability of non-native species to establish in native communities with acquisitive leaf strategies (LaForgia et al., 2020; Lemoine et al., 2015; Li et al., 2022; Lodge et al., 2018). In other words, our results suggest that in mountain ecosystems conservative communities may be inherently less hospitable to fast-growing, acquisitive invaders that usually rely on high resource availability. Beyond trait dominance, we also detected an effect of functional diversity on invasion dynamics, which was positively related to non-native species abundance but not to their occurrence. This finding suggests that functional diversity in native communities does not necessarily prevent the establishment of non-native species but may influence the extent to which they can proliferate once present. A possible explanation lies in the greater heterogeneity of strategies within functionally diverse communities, which may create a broader range of ecological niches and microhabitats that can be exploited by invaders (Davies et al., 2005; Fridley et al., 2007). In this context, higher functional diversity may reduce competitive exclusion and facilitate coexistence, thereby allowing non-native species to increase in cover. Alternatively, functionally diverse communities may often reflect intermediate successional stages or more dynamic environments, where resource turnover and disturbance create windows of opportunity for invasion (Funk et al., 2008; Shea & Chesson, 2002). The fact that functional diversity affected abundance but not occurrence supports the idea that climate and disturbance remain the primary filters controlling whether non-native plants can establish, while biotic interactions and community structure become more important in determining their success after establishment.

### Biotic resistance modulates the effect of global changes on plant invasion

As global changes are expected to reshuffle not only non-native species but also natives (Lembrechts et al., 2017a; Pauchard et al., 2016; Petitpierre et al., 2016), this study aims to examine how global changes altered the resistance of native communities to invasion. Our findings show that road disturbance and changes occurred in land-use and climate shaped the composition and structure of native communities and, in turn, invasion dynamics. In particular, we found that road disturbance increased the dominance of acquisitive, fast-growing species, which thrive in disturbed environments where competition for resources is reduced, and environmental conditions are more favorable to opportunistic growth strategies (Alvarez et al., 2025; Herben et al., 2018). These native plant communities offer favorable conditions for non-native plants establishment. Therefore, roads do not only favor invasion by acting as dispersal corridors and propagule reservoirs (Lembrechts et al., 2017a), but also by changing native community composition.

Land-use changes occurred over the last 70 years in the study area also contributed to alter native plants community composition, influencing invasion dynamics indirectly. In many European mountains, landscapes historically dominated by pastures and agricultural lands have undergone significant changes due to widespread agricultural abandonment and rural exodus since World War II (Lasanta et al., 2017). This shift has triggered natural vegetation regrowth and over a few decades these areas have experienced substantial forest recovery (Bracchetti et al., 2012; Sitzia et al., 2010). These succession processes have been observed also in the study area (Evangelista et al., 2016; Frate et al., 2018; Malavasi et al., 2018) and deeply transformed the vegetation structure by increasing conservative communities, with important implications for native communities’ ability to resist plant invasions. Our findings show that forest expansion decreased the dominance of acquisitive species in native communities which in turn become less favorable to the establishment and expansion of non-native species. This is because, unlike acquisitive communities. conservative ones are better equipped for resource conservation and potentially create stable conditions that reduce niche opportunities for invaders. In line with our results, other studies already illustrated that mountain forests are still relatively resistant to invasion (Catford et al., 2012; Von Holle et al., 2003; Wagner et al., 2017). The dense canopy cover, understory, and thick litter layer of a forest could limit the ability of light-demanding, fast-growing invaders to succeed, thus creating physical barriers to invasion. Conversely, we also found that in sites where agricultural abandonment in the surrounding resulted in transitional or early successional stages, such as meadows, the reduced structural complexity and lower plant size of the native vegetation have created opportunities for non-native species to establish. Our study thus suggests that in Mediterranean mountain ecosystems, the gradual abandonment of agricultural lands and subsequent forest expansion are progressively enhancing the biotic resistance of native communities, whereas the low- to middle elevations grasslands maintain a higher vulnerability to plant invasion. Supporting this, we observed that out of the 16 non-native species identified in our study, only five were tree species, and they predominantly invaded roadside plots rather than penetrating forest interiors (Santoianni et al., 2024, 2025). However, the most abundant non-native species in our study area was *Ailanthus altissima* (Mill.) S an invasive tree species, raising important considerations. *A. altissima* has been recognized as a highly adaptable and aggressive invader, capable of exploiting disturbed habitats and encroaching into less disturbed forests (Kowarik & Säumel, 2007; Marzialetti et al., 2025; Sladonja et al., 2015). Even though, according to our results, native forests currently exhibit high resistance to invasion and climate change has not yet impacted the composition of native communities, these dynamics could shift if climatic conditions worsen or other global change factors, such as human disturbance, weaken native community resilience. In such scenarios, invasive species like *A*. *altissima* may seize the opportunity to expand further into forested habitats (Cao Pinna et al., 2024; Essl et al., 2012).

## ACKNOWLEDGEMENTS

We would like to thank the Mountain Invasion Research Network (MIREN) consortium for methodological planning; the Maiella National Park staff and Gran Sasso and Monti della Laga National Park (GSMLNP) staff for logistic and technical support; Marco Andrello, Marco Varricchione, Simona Sarmati, Viola Filippini, Alessandro Zerbini, Maurizio Cutini, and Silvia Giulio for their assistance in collecting data in the field.

## AUTHOR CONTRIBUTIONS

MC and GLB conceived and designed the study. All authors collected the data. GLB and MC analysed and interpreted the data. GLB wrote the first draft of the manuscript. All authors made significant contributions to the manuscript and approved it for publication.

## DATA AVAILABILITY STATEMENT

The data that support the findings of this study are available in Santoianni et al. (2025).

## REFERENCES

Alexander, J. M., Chalmandrier, L., Lenoir, J., Burgess, T. I., Essl, F., Haider, S., Kueffer, C., McDougall, K., Milbau, A., Nuñez, M. A., Pauchard, A., Rabitsch, W., Rew, L. J., Sanders, N. J., & Pellissier, L. (2018). Lags in the response of mountain plant communities to climate change. Global Change Biology, 24(2), 563–579. 10.1111/gcb.13976

Alexander, J. M., Diez, J. M., & Levine, J. M. (2015). Novel competitors shape species’ responses to climate change. Nature, 525(7570), 515–518. 10.1038/nature14952

Alexander, J. M., Lembrechts, J. J., Cavieres, L. A., Daehler, C., Haider, S., Kueffer, C., Liu, G., McDougall, K., Milbau, A., Pauchard, A., Rew, L. J., & Seipel, T. (2016). Plant invasions into mountains and alpine ecosystems: Current status and future challenges. Alpine Botany, 126(2), 89–103. 10.1007/s00035-016-0172-8

Alvarez, M. A., Cavieres, L. A., Aschero, V., Bonjour, L. D. J., & Barros, A. (2025). Road Disturbance Shapes the Functional Composition of Native Plant Species but Not That of Non-Natives in the Arid Andes. Applied Vegetation Science, 28(2), e70031. 10.1111/avsc.70031

Barros, A., Fuentes Lillo, E., Aschero, V., Pauchard, A., Alvarez, M. A., Wedegärtner, R., Clavel, J., Müllerová, J., Pergl, J., Zong, S., Vítková, M., Klinerová, T., Cavieres, L. A., Larson, C., Rew, L. J., Seipel, T., Meffre, C., Arellano, T., Essl, F., … Lembrechts, J. J. (2025). Beyond the Trail—Understanding Non-Native Plant Invasions in Mountain Ecosystems. Global Ecology and Biogeography, 34(6), e70060. 10.1111/geb.70060

Bartolucci, F., Peruzzi, L., Galasso, G., Alessandrini, A., Ardenghi, N. M. G., Bacchetta, G., Banfi, E., Barberis, G., Bernardo, L., Bouvet, D., Bovio, M., Calvia, G., Castello, M., Cecchi, L., Del Guacchio, E., Domina, G., Fascetti, S., Gallo, L., Gottschlich, G., … Conti, F. (2024). A second update to the checklist of the vascular flora native to Italy. Plant Biosystems - An International Journal Dealing with All Aspects of Plant Biology, 158(2), 219–296. 10.1080/11263504.2024.2320126

Bates, D., Mächler, M., Bolker, B., & Walker, S. (2014). Fitting Linear Mixed-Effects Models using lme4. 10.48550/ARXIV.1406.5823

Bjorkman, A. D., Myers-Smith, I. H., Elmendorf, S. C., Normand, S., Rüger, N., Beck, P. S. A., Blach-Overgaard, A., Blok, D., Cornelissen, J. H. C., Forbes, B. C., Georges, D., Goetz, S. J., Guay, K. C., Henry, G. H. R., HilleRisLambers, J., Hollister, R. D., Karger, D. N., Kattge, J., Manning, P., … Weiher, E. (2018). Plant functional trait change across a warming tundra biome. Nature, 562(7725), 57–62. 10.1038/s41586-018-0563-7

Bracchetti, L., Carotenuto, L., & Catorci, A. (2012). Land-cover changes in a remote area of central Apennines (Italy) and management directions. Landscape and Urban Planning, 104(2), 157–170. 10.1016/j.landurbplan.2011.09.005

Burnham, K. P., & Anderson, D. R. (2002). Model selection and multimodel inference: A practical information-theoretic approach. 2nd Ed. New York, Springer-Verlag.

Cao Pinna, L., Axmanová, I., Chytrý, M., Malavasi, M., Acosta, A. T. R., Giulio, S., Attorre, F., Bergmeier, E., Biurrun, I., Campos, J. A., Font, X., Küzmič, F., Landucci, F., Marcenò, C., Rodríguez-Rojo, M. P., & Carboni, M. (2021). The biogeography of alien plant invasions in the Mediterranean Basin. Journal of Vegetation Science, 32(2), e12980. 10.1111/jvs.12980

Cao Pinna, L., Gallien, L., Pollock, L. J., Axmanová, I., Chytrý, M., Malavasi, M., Acosta, A. T. R., Antonio Campos, J., & Carboni, M. (2024). Plant invasion in Mediterranean Europe: Current hotspots and future scenarios. Ecography, 2024(5), e07085. 10.1111/ecog.07085

Carboni, M., Calderon-Sanou, I., Pollock, L., Violle, C., DivGrass Consortium, & Thuiller, W. (2018). Functional traits modulate the response of alien plants along abiotic and biotic gradients. Global Ecology and Biogeography, 27(10), 1173–1185. 10.1111/geb.12775

Carboni, M., Guéguen, M., Barros, C., Georges, D., Boulangeat, I., Douzet, R., Dullinger, S., Klonner, G., Van Kleunen, M., Essl, F., Bossdorf, O., Haeuser, E., Talluto, M. V., Moser, D., Block, S., Conti, L., Dullinger, I., Münkemüller, T., & Thuiller, W. (2018). Simulating plant invasion dynamics in mountain ecosystems under global change scenarios. Global Change Biology, 24(1). 10.1111/gcb.13879

Carboni, M., Münkemüller, T., Gallien, L., Lavergne, S., Acosta, A., & Thuiller, W. (2013). Darwin’s naturalization hypothesis: Scale matters in coastal plant communities. Ecography, 36(5), 560–568. 10.1111/j.1600-0587.2012.07479.x

Carboni, M., Münkemüller, T., Lavergne, S., Choler, P., Borgy, B., Violle, C., Essl, F., Roquet, C., Munoz, F., DivGrass Consortium, & Thuiller, W. (2016). What it takes to invade grassland ecosystems: Traits, introduction history and filtering processes. Ecology Letters, 19(3), 219–229. 10.1111/ele.12556

Catford, J. A., Vesk, P. A., Richardson, D. M., & Pyšek, P. (2012). Quantifying levels of biological invasion: Towards the objective classification of invaded and invasible ecosystems. Global Change Biology, 18(1), 44–62. 10.1111/j.1365-2486.2011.02549.x

Cavieres, L. A. (2021). Facilitation and the invasibility of plant communities. Journal of Ecology, 109(5), 2019–2028. 10.1111/1365-2745.13627

Christen, D. C., & Matlack, G. R. (2009). The habitat and conduit functions of roads in the spread of three invasive plant species. Biological Invasions, 11(2), 453–465. 10.1007/s10530-008-9262-x

Collins, C. G., Elmendorf, S. C., Smith, J. G., Shoemaker, L., Szojka, M., Swift, M., & Suding, K. N. (2022). Global change re-structures alpine plant communities through interacting abiotic and biotic effects. Ecology Letters, 25(8), 1813–1826. 10.1111/ele.14060

Congalton, R. G., & Green, K. (2019). Assessing the Accuracy of Remotely Sensed Data: Principles and Practices, Third Edition (3rd ed.). CRC Press. 10.1201/9780429052729

Conti, L., Block, S., Parepa, M., Münkemüller, T., Thuiller, W., Acosta, A. T. R., Van Kleunen, M., Dullinger, S., Essl, F., Dullinger, I., Moser, D., Klonner, G., Bossdorf, O., & Carboni, M. (2018). Functional trait differences and trait plasticity mediate biotic resistance to potential plant invaders. Journal of Ecology, 106(4), 1607–1620. 10.1111/1365-2745.12928

Cutini, M., Flavio, M., Giuliana, B., Guido, R., & Jean-Paul, T. (2021). Bioclimatic pattern in a Mediterranean mountain area: Assessment from a classification approach on a regional scale. International Journal of Biometeorology, 65(7), 1085–1097. 10.1007/s00484-021-02089-x

Daehler, C. C. (2001). Darwin’s Naturalization Hypothesis Revisited. The American Naturalist, 158(3), 324–330. 10.1086/321316

Dainese, M., Aikio, S., Hulme, P. E., Bertolli, A., Prosser, F., & Marini, L. (2017). Human disturbance and upward expansion of plants in a warming climate. Nature Climate Change, 7(8), Article 8. 10.1038/nclimate3337

Dainese, M., Crepaz, H., Bottarin, R., Fontana, V., Guariento, E., Hilpold, A., Obojes, N., Paniccia, C., Scotti, A., Seeber, J., Steinwandter, M., Tappeiner, U., & Niedrist, G. (2024). Global change experiments in mountain ecosystems: A systematic review. Ecological Monographs, 94(4), e1632. 10.1002/ecm.1632

Davies, K. F., Chesson, P., Harrison, S., Inouye, B. D., Melbourne, B. A., & Rice, K. J. (2005). Spatial heterogeneity explains the scale dependence of the native–exotic diversity relationship. Ecology, 86(6), 1602–1610. 10.1890/04-1196

Dawson, W., Rohr, R. P., Van Kleunen, M., & Fischer, M. (2012). Alien plant species with a wider global distribution are better able to capitalize on increased resource availability. New Phytologist, 194(3), 859–867. 10.1111/j.1469-8137.2012.04104.x

de Bello, F., Botta-Dukát, Z., Lepš, J., & Fibich, P. (2021). Towards a more balanced combination of multiple traits when computing functional differences between species. Methods in Ecology and Evolution, 12(3), 443–448.

de Bello, F., P. Carmona, C., Leps, J., Szava-Kovats, R., & Pärtel, M. (2016). Functional diversity through the mean trait dissimilarity: Resolving shortcomings with existing paradigms and algorithms. Oecologia, 180. 10.1007/s00442-016-3546-0

Díaz, S., Kattge, J., Cornelissen, J. H. C., Wright, I. J., Lavorel, S., Dray, S., Reu, B., Kleyer, M., Wirth, C., Colin Prentice, I., Garnier, E., Bönisch, G., Westoby, M., Poorter, H., Reich, P. B., Moles, A. T., Dickie, J., Gillison, A. N., Zanne, A. E., … Gorné, L. D. (2016). The global spectrum of plant form and function. Nature, 529(7585), Article 7585. 10.1038/nature16489

Drenovsky, R. E., & James, J. J. (2010). Designing Invasion-Resistant Plant Communities: The Role of Plant Functional Traits. Rangelands, 32(1), 32–37. 10.2111/RANGELANDS-D-09-00002.1

Duncan, R. P., & Williams, P. A. (2002). Darwin’s naturalization hypothesis challenged. Nature, 417(6889), 608–609. 10.1038/417608a

Elton, C. S. (1958). The ecology of invasions by animals and plants. London: Methuen.

Emery, S. M., & Gross, K. L. (2007). Dominant species identity, not community evenness, regulates invasion in experimental grassland plant communities. Ecology, 88(4), 954–964.

Essl, F., Dullinger, S., Rabitsch, W., Hulme, P. E., Pyšek, P., Wilson, J. R. U., & Richardson, D. M. (2015). Historical legacies accumulate to shape future biodiversity in an era of rapid global change. Diversity and Distributions, 21(5), 534–547. 10.1111/ddi.12312

Essl, F., Mang, T., & Moser, D. (2012). Ancient and recent alien species in temperate forests: Steady state and time lags. Biological Invasions, 14(7), 1331–1342. 10.1007/s10530-011-0156-y

Evangelista, A., Frate, L., Carranza, M. L., Attorre, F., Pelino, G., & Stanisci, A. (2016). Changes in composition, ecology and structure of high-mountain vegetation: A re-visitation study over 42 years. AoB PLANTS, 8, plw004. 10.1093/aobpla/plw004

Fan, S., Yang, Q., Li, S., Fristoe, T. S., Cadotte, M. W., Essl, F., Kreft, H., Pergl, J., Pyšek, P., Weigelt, P., Kartesz, J., Nishino, M., Wieringa, J. J., & Van Kleunen, M. (2023). A latitudinal gradient in Darwin’s naturalization conundrum at the global scale for flowering plants. Nature Communications, 14(1), 6244. 10.1038/s41467-023-41607-w

Frate, L., Carranza, M. L., Evangelista, A., Stinca, A., Schaminée, J. H. J., & Stanisci, A. (2018). Climate and land use change impacts on Mediterranean high-mountain vegetation in the Apennines since the 1950s. Plant Ecology & Diversity, 11(1), 85–96. 10.1080/17550874.2018.1473521

Fridley, J. D., Stachowicz, J. J., Naeem, S., Sax, D. F., Seabloom, E. W., Smith, M. D., Stohlgren, T. J., Tilman, D., & Holle, B. V. (2007). The invasion paradox: Reconciling pattern and process in species invasions. Ecology, 88(1), 3–17. 10.1890/0012-9658(2007)88[3:TIPRPA]2.0.CO;2

Fuentes-Lillo, E., Lembrechts, J. J., Cavieres, L. A., Jiménez, A., Haider, S., Barros, A., & Pauchard, A. (2021). Anthropogenic factors overrule local abiotic variables in determining non-native plant invasions in mountains. Biological Invasions, 23(12), 3671–3686. 10.1007/s10530-021-02602-8

Funk, J. L., Cleland, E. E., Suding, K. N., & Zavaleta, E. S. (2008). Restoration through reassembly: Plant traits and invasion resistance. Trends in Ecology & Evolution, 23(12), 695–703.

Galasso, G., Conti, F., Peruzzi, L., Alessandrini, A., Ardenghi, N. M. G., Bacchetta, G., Banfi, E., Barberis, G., Bernardo, L., Bouvet, D., Bovio, M., Castello, M., Cecchi, L., Del Guacchio, E., Domina, G., Fascetti, S., Gallo, L., Guarino, R., Gubellini, L., … Bartolucci, F. (2024). A second update to the checklist of the vascular flora alien to Italy. Plant Biosystems - An International Journal Dealing with All Aspects of Plant Biology, 158(2), 297–340. 10.1080/11263504.2024.2320129

Galasso, G., Conti, F., Peruzzi, L., Ardenghi, N. M. G., Banfi, E., Celesti-Grapow, L., Albano, A., Alessandrini, A., Bacchetta, G., Ballelli, S., Bandini Mazzanti, M., Barberis, G., Bernardo, L., Blasi, C., Bouvet, D., Bovio, M., Cecchi, L., Del Guacchio, E., Domina, G., … Bartolucci, F. (2018). An updated checklist of the vascular flora alien to Italy. Plant Biosystems - An International Journal Dealing with All Aspects of Plant Biology, 152(3), 556–592. 10.1080/11263504.2018.1441197

Gallien, L., & Carboni, M. (2017). The community ecology of invasive species: Where are we and what’s next? Ecography, 40(2), 335–352. 10.1111/ecog.02446

Gallien, L., Mazel, F., Lavergne, S., Renaud, J., Douzet, R., & Thuiller, W. (2015). Contrasting the effects of environment, dispersal and biotic interactions to explain the distribution of invasive plants in alpine communities. Biological Invasions, 17(5), 1407–1423. 10.1007/s10530-014-0803-1

Gana, K., & Broc, G. (2019). Structural equation modeling with lavaan. John Wiley & Sons.

Geppert, C., Boscutti, F., La Bella, G., Marchi, V., Corcos, D., Filippi, A., & Marini, L. (2021). Contrasting response of native and non-native plants to disturbance and herbivory in mountain environments. Journal of Biogeography, 1–12. 10.1111/jbi.14097

Gottfried, M., Pauli, H., Futschik, A., Akhalkatsi, M., Barančok, P., Benito Alonso, J. L., Coldea, G., Dick, J., Erschbamer, B., Fernández Calzado, M. R., Kazakis, G., Krajči, J., Larsson, P., Mallaun, M., Michelsen, O., Moiseev, D., Moiseev, P., Molau, U., Merzouki, A., … Grabherr, G. (2012). Continent-wide response of mountain vegetation to climate change. Nature Climate Change, 2(2), 111–115. 10.1038/nclimate1329

Gu, Z., Gu, M. Z., & GlobalOptions, I. (2024). Package ‘circlize’.

Haider, S., Kueffer, C., Bruelheide, H., Seipel, T., Alexander, J. M., Rew, L. J., Arévalo, J. R., Cavieres, L. A., McDougall, K. L., Milbau, A., Naylor, B. J., Speziale, K., & Pauchard, A. (2018). Mountain roads and non-native species modify elevational patterns of plant diversity. Global Ecology and Biogeography, 27(6), 667–678. 10.1111/geb.12727

Haider, S., Lembrechts, J. J., McDougall, K., Pauchard, A., Alexander, J. M., Barros, A., Cavieres, L. A., Rashid, I., Rew, L. J., Aleksanyan, A., Arévalo, J. R., Aschero, V., Chisholm, C., Clark, V. R., Clavel, J., Daehler, C., Dar, P. A., Dietz, H., Dimarco, R. D., … Seipel, T. (2022). Think globally, measure locally: The MIREN standardized protocol for monitoring plant species distributions along elevation gradients. Ecology and Evolution, 12(2), e8590. 10.1002/ece3.8590

Heberling, J. M., & Fridley, J. D. (2013). Resource-use strategies of native and invasive plants in Eastern North American forests. New Phytologist, 200(2), 523–533. 10.1111/nph.12388

Herben, T., Klimešová, J., & Chytrý, M. (2018). Effects of disturbance frequency and severity on plant traits: An assessment across a temperate flora. Functional Ecology, 32(3), 799–808. 10.1111/1365-2435.13011

Hesselbarth, M. H. K., Sciaini, M., Nowosad, J., & Hanss, S. (2018). landscapemetrics: Landscape Metrics for Categorical Map Patterns (p. 2.1.4) [Dataset]. 10.32614/CRAN.package.landscapemetrics

Hoch, G., & Körner, C. (2012). Global patterns of mobile carbon stores in trees at the high-elevation tree line. Global Ecology and Biogeography, 21(8), 861–871. 10.1111/j.1466-8238.2011.00731.x

Hoffman, G. E., & Schadt, E. E. (2016). variancePartition: Interpreting drivers of variation in complex gene expression studies. BMC Bioinformatics, 17(1), 483. 10.1186/s12859-016-1323-z

Ibáñez, I., Liu, G., Petri, L., Schaffer-Morrison, S., & Schueller, S. (2021). Assessing vulnerability and resistance to plant invasions: A native community perspective. Invasive Plant Science and Management, 14(2), 64–74. 10.1017/inp.2021.15

Karger, D. N., & Zimmermann, N. E. (2018). CHELSAcruts—High resolution temperature and precipitation timeseries for the 20th century and beyond (Version 1.0) [Geotiff,PDF]. EnviDat. 10.16904/ENVIDAT.159

Kattge, J., Bönisch, G., Díaz, S., Lavorel, S., Prentice, I. C., Leadley, P., Tautenhahn, S., Werner, G. D., Aakala, T., & Abedi, M. (2020). TRY plant trait database–enhanced coverage and open access. Global Change Biology, 26(1), 119–188.

Körner, C. (2007). The use of ‘altitude’ in ecological research. Trends in Ecology & Evolution, 22(11), 569–574. 10.1016/j.tree.2007.09.006

Kowarik, I., & Säumel, I. (2007). Biological flora of Central Europe: Ailanthus altissima (Mill.) Swingle. Perspectives in Plant Ecology, Evolution and Systematics, 8(4), 207–237. 10.1016/j.ppees.2007.03.002

Kraft, N. J. B., Godoy, O., & Levine, J. M. (2015). Plant functional traits and the multidimensional nature of species coexistence. Proceedings of the National Academy of Sciences, 112(3), 797–802. 10.1073/pnas.1413650112

LaForgia, M. L., Harrison, S. P., & Latimer, A. M. (2020). Invasive species interact with climatic variability to reduce success of natives. Ecology, 101(6), e03022. 10.1002/ecy.3022

Laliberté, E., Legendre, P., Shipley, B., & Laliberté, M. E. (2014). Package ‘FD’. Measuring Functional Diversity from Multiple Traits, and Other Tools for Functional Ecology, 1.0-12.

Lasanta, T., Arnáez, J., Pascual, N., Ruiz-Flaño, P., Errea, M. P., & Lana-Renault, N. (2017). Space–time process and drivers of land abandonment in Europe. CATENA, 149, 810– 823. 10.1016/j.catena.2016.02.024

Le, H., Mao, J., Cavender-Bares, J., Pinto-Ledezma, J. N., Deng, Y., Zhao, C., Xiong, G., Xu, W., & Xie, Z. (2024). Non-native plants tend to be phylogenetically distant but functionally similar to native plants under intense disturbance at the Three Gorges Reservoir Area. New Phytologist, 244(5), 2078–2088. 10.1111/nph.20126

Lefcheck, J. S. (2016). piecewiseSEM: Piecewise structural equation modelling in r for ecology, evolution, and systematics. Methods in Ecology and Evolution, 7(5), 573–579. 10.1111/2041-210X.12512

Lejeune, R., Fuentes-Lillo, E., Haesen, S., Pirée, A., Wiegmans, D., Hostens, L., Lenoir, J., Pergl, J., Vítková, M., Seipel, T., Kutlvašr, J., Nuñez, M. A., Dimarco, R. D., Alexander, J., Backes, A. R., Haider, S., Pauchard, A., Nijs, I., & Lembrechts, J. J. (2024). Mountain roads across the globe significantly alter local soil microclimates. 10.1101/2024.11.28.625797

Lembrechts, J. J., Alexander, J. M., Cavieres, L. A., Haider, S., Lenoir, J., Kueffer, C., McDougall, K., Naylor, B. J., Nuñez, M. A., Pauchard, A., Rew, L. J., Nijs, I., & Milbau, A. (2017a). Mountain roads shift native and non-native plant species’ ranges. Ecography, 40(3), 353–364. 10.1111/ecog.02200

Lembrechts, J. J., Alexander, J. M., Cavieres, L. A., Haider, S., Lenoir, J., Kueffer, C., McDougall, K., Naylor, B. J., Nuñez, M. A., Pauchard, A., Rew, L. J., Nijs, I., & Milbau, A. (2017b). Mountain roads shift native and non-native plant species’ ranges. Ecography, 40(3), 353–364. 10.1111/ecog.02200

Lembrechts, J. J., Pauchard, A., Lenoir, J., Nuñez, M. A., Geron, C., Ven, A., Bravo-Monasterio, P., Teneb, E., Nijs, I., & Milbau, A. (2016). Disturbance is the key to plant invasions in cold environments. Proceedings of the National Academy of Sciences, 113(49), 14061–14066. 10.1073/pnas.1608980113

Lemoine, N. P., Shue, J., Verrico, B., Erickson, D., Kress, W. J., & Parker, J. D. (2015). Phylogenetic relatedness and leaf functional traits, not introduced status, influence community assembly. Ecology, 96(10), 2605–2612. 10.1890/14-1883.1

Lenoir, J., Gégout, J. C., Marquet, P. A., De Ruffray, P., & Brisse, H. (2008). A Significant Upward Shift in Plant Species Optimum Elevation During the 20th Century. Science, 320(5884), 1768–1771. 10.1126/science.1156831

Li, S., Jia, P., Fan, S., Wu, Y., Liu, X., Meng, Y., Li, Y., Shu, W., Li, J., & Jiang, L. (2022). Functional traits explain the consistent resistance of biodiversity to plant invasion under nitrogen enrichment. Ecology Letters, 25(4), 778–789. 10.1111/ele.13951

Lodge, A. G., Whitfeld, T. J. S., Roth, A. M., & Reich, P. B. (2018). Invasive plants in Minnesota are “joining the locals”: A trait-based analysis. Journal of Vegetation Science, 29(4), 746–755. 10.1111/jvs.12659

MacArthur, R., & Levins, R. (1967). The limiting similarity, convergence, and divergence of coexisting species. The American Naturalist, 101(921), 377–385.

Malavasi, M., Carranza, M. L., Moravec, D., & Cutini, M. (2018). Reforestation dynamics after land abandonment: A trajectory analysis in Mediterranean mountain landscapes. Regional Environmental Change, 18(8), 2459–2469. 10.1007/s10113-018-1368-9

Malavasi, M., Santoro, R., Cutini, M., Acosta, A. T. R., & Carranza, M. L. (2013). What has happened to coastal dunes in the last half century? A multitemporal coastal landscape analysis in Central Italy. Landscape and Urban Planning, 119, 54–63. 10.1016/j.landurbplan.2013.06.012

Marzialetti, F., Lozano, V., Große-Stoltenberg, A., Carranza, M. L., Innangi, M., La Bella, G., Bagella, S., Rivieccio, G., Bacchetta, G., Podda, L., & Brundu, G. (2025). Assessing eco-physiological patterns of Ailanthus altissima (Mill.) Swingle and differences with native vegetation using Copernicus satellite data on a Mediterranean Island. Ecological Informatics, 87, 103080. 10.1016/j.ecoinf.2025.103080

McDougall, K. L., Lembrechts, J., Rew, L. J., Haider, S., Cavieres, L. A., Kueffer, C., Milbau, A., Naylor, B. J., Nuñez, M. A., Pauchard, A., Seipel, T., Speziale, K. L., Wright, G. T., & Alexander, J. M. (2018). Running off the road: Roadside non-native plants invading mountain vegetation. Biological Invasions, 20(12), 3461–3473. 10.1007/s10530-018-1787-z

Mengist, W., Soromessa, T., & Legese, G. (2020). Ecosystem services research in mountainous regions: A systematic literature review on current knowledge and research gaps. Science of The Total Environment, 702, 134581. 10.1016/j.scitotenv.2019.134581

Milbau, A., Shevtsova, A., Osler, N., Mooshammer, M., & Graae, B. J. (2013). Plant community type and small-scale disturbances, but not altitude, influence the invasibility in subarctic ecosystems. New Phytologist, 197(3), 1002–1011. 10.1111/nph.12054

Naimi, B. (2012). usdm: Uncertainty Analysis for Species Distribution Models (p. 2.1-7) [Dataset]. 10.32614/CRAN.package.usdm

Oksanen, J., Blanchet, F. G., Friendly, M., Kindt, R., Legendre, P., McGlinn, D., Minchin, P. R., O’Hara, R. B., Simpson, G. L., & Solymos, P. (2021). Vegan: Community Ecology Package. R package version 2.5-7. 2020.

Pakeman, R. J., & Quested, H. M. (2007). Sampling plant functional traits: What proportion of the species need to be measured? Applied Vegetation Science, 10(1), 91–96.

Park, D. S., Feng, X., Maitner, B. S., Ernst, K. C., & Enquist, B. J. (2020). Darwin’s naturalization conundrum can be explained by spatial scale. Proceedings of the National Academy of Sciences, 117(20), 10904–10910. 10.1073/pnas.1918100117

Pauchard, A., & Alaback, P. B. (2004). Influence of Elevation, Land Use, and Landscape Context on Patterns of Alien Plant Invasions along Roadsides in Protected Areas of South-Central Chile. Conservation Biology, 18(1), 238–248. 10.1111/j.1523-1739.2004.00300.x

Pauchard, A., Kueffer, C., Dietz, H., Daehler, C. C., Alexander, J., Edwards, P. J., Arévalo, J. R., Cavieres, L. A., Guisan, A., Haider, S., Jakobs, G., McDougall, K., Millar, C. I., Naylor, B. J., Parks, C. G., Rew, L. J., & Seipel, T. (2009). Ain’t no mountain high enough: Plant invasions reaching new elevations. Frontiers in Ecology and the Environment, 7(9), 479–486. 10.1890/080072

Pauchard, A., Milbau, A., Albihn, A., Alexander, J., Burgess, T., Daehler, C., Englund, G., Essl, F., Evengård, B., Greenwood, G. B., Haider, S., Lenoir, J., McDougall, K., Muths, E., Nuñez, M. A., Olofsson, J., Pellissier, L., Rabitsch, W., Rew, L. J., … Kueffer, C. (2016). Non-native and native organisms moving into high elevation and high latitude ecosystems in an era of climate change: New challenges for ecology and conservation. Biological Invasions, 18(2), 345–353. 10.1007/s10530-015-1025-x

Pesaresi, S., Biondi, E., & Casavecchia, S. (2017). Bioclimates of Italy. Journal of Maps, 13(2), 955–960. 10.1080/17445647.2017.1413017

Petitpierre, B., McDougall, K., Seipel, T., Broennimann, O., Guisan, A., & Kueffer, C. (2016). Will climate change increase the risk of plant invasions into mountains? Ecological Applications, 26(2), 530–544.

Pyšek, P., Hulme, P. E., Simberloff, D., Bacher, S., Blackburn, T. M., Carlton, J. T., Dawson, W., Essl, F., Foxcroft, L. C., Genovesi, P., Jeschke, J. M., Kühn, I., Liebhold, A. M., Mandrak, N. E., Meyerson, L. A., Pauchard, A., Pergl, J., Roy, H. E., Seebens, H., … Richardson, D. M. (2020). Scientists’ warning on invasive alien species. Biological Reviews, 95(6), 1511–1534. 10.1111/brv.12627

R Core Team. (2023). R: A Language and Environment for Statistical Computing. R Foundation for Statistical Computing. https://www.R-project.org/

Reich, P. B. (2014). The world-wide ‘fast-slow’ plant economics spectrum: A traits manifesto. Journal of Ecology, 102(2), 275–301. 10.1111/1365-2745.12211

Richards, S. A. (2005). Testing ecological theory using the information-theoretic approach: Examples and cautionary results. Ecology, 86(10), 2805–2814. 10.1890/05-0074

Richardson, D. M., & Pyšek, P. (2006). Plant invasions: Merging the concepts of species invasiveness and community invasibility. Progress in Physical Geography: Earth and Environment, 30(3), 409–431. 10.1191/0309133306pp490pr

Rodewald, A. D., & Arcese, P. (2016). Direct and Indirect Interactions between Landscape Structure and Invasive or Overabundant Species. Current Landscape Ecology Reports, 1(1), 30–39. 10.1007/s40823-016-0004-y

Roy, H. E., Pauchard, A., Stoett, P., Renard Truong, T., Bacher, S., Galil, B. S., Hulme, P. E., Ikeda, T., Sankaran, K., McGeoch, M. A., Meyerson, L. A., Nuñez, M. A., Ordonez, A., Rahlao, S. J., Schwindt, E., Seebens, H., Sheppard, A. W., & Vandvik, V. (2024). IPBES Invasive Alien Species Assessment: Summary for Policymakers (Version 3). Zenodo. 10.5281/ZENODO.7430692

Santoianni, L. A., Bartolucci, F., Carboni, M., Conti, F., La Bella, G., Varricchione, M., & Stanisci, A. (2025). MARA Vegetation Database: Monitoring Alien species along mountain Roads in the central Apennines. Vegetation Ecology and Diversity, 62, 1–9. 10.3897/ved.139363

Santoianni, L. A., Innangi, M., Varricchione, M., Carboni, M., La Bella, G., Haider, S., & Stanisci, A. (2024). Ecological features facilitating spread of alien plants along Mediterranean mountain roads. Biological Invasions, 26(11), 3879–3899. 10.1007/s10530-024-03418-y

Schirpke, U., Wang, G., & Padoa-Schioppa, E. (2021). Editorial: Mountain landscapes: Protected areas, ecosystem services, and future challenges. Ecosystem Services, 49, 101302. 10.1016/j.ecoser.2021.101302

Seebens, H., Blackburn, T. M., Dyer, E. E., Genovesi, P., Hulme, P. E., Jeschke, J. M., Pagad, S., Pyšek, P., Winter, M., Arianoutsou, M., Bacher, S., Blasius, B., Brundu, G., Capinha, C., Celesti-Grapow, L., Dawson, W., Dullinger, S., Fuentes, N., Jäger, H., … Essl, F. (2017). No saturation in the accumulation of alien species worldwide. Nature Communications, 8(1), 14435. 10.1038/ncomms14435

Seebens, H., Blackburn, T. M., Hulme, P. E., Van Kleunen, M., Liebhold, A. M., Orlova-Bienkowskaja, M., Pyšek, P., Schindler, S., & Essl, F. (2021). Around the world in 500 years: Inter-regional spread of alien species over recent centuries. Global Ecology and Biogeography, 30(8), 1621–1632. 10.1111/geb.13325

Seipel, T., Alexander, J. M., Edwards, P. J., & Kueffer, C. (2016). Range limits and population dynamics of non-native plants spreading along elevation gradients. Perspectives in Plant Ecology, Evolution and Systematics, 20, 46–55. 10.1016/j.ppees.2016.04.001

Shea, K., & Chesson, P. (2002). Community ecology theory as a framework for biological invasions. Trends in Ecology & Evolution, 17(4), 170–176. 10.1016/S0169-5347(02)02495-3

Sheppard, C. S. (2019). Relative performance of co-occurring alien plant invaders depends on traits related to competitive ability more than niche differences. Biological Invasions, 21(4), 1101–1114. 10.1007/s10530-018-1884-z

Shipley, B. (2013). The AIC model selection method applied to path analytic models compared using ad-separation test. Ecology, 94(3), 560–564.

Simpson, K. J., Atkinson, R. R. L., Mockford, E. J., Bennett, C., Osborne, C. P., & Rees, M. (2021). Large seeds provide an intrinsic growth advantage that depends on leaf traits and root allocation. Functional Ecology, 35(10), 2168–2178. 10.1111/1365-2435.13871

Sitzia, T., Semenzato, P., & Trentanovi, G. (2010). Natural reforestation is changing spatial patterns of rural mountain and hill landscapes: A global overview. Forest Ecology and Management, 259(8), 1354–1362. 10.1016/j.foreco.2010.01.048

Sladonja, B., Sušek, M., & Guillermic, J. (2015). Review on Invasive Tree of Heaven (Ailanthus altissima (Mill.) Swingle) Conflicting Values: Assessment of Its Ecosystem Services and Potential Biological Threat. Environmental Management, 56(4), 1009– 1034. 10.1007/s00267-015-0546-5

Steinbauer, K., Lamprecht, A., Winkler, M., Di Cecco, V., Fasching, V., Ghosn, D., Maringer, A., Remoundou, I., Suen, M., Stanisci, A., Venn, S., & Pauli, H. (2022). Recent changes in high-mountain plant community functional composition in contrasting climate regimes. Science of The Total Environment, 829, 154541. 10.1016/j.scitotenv.2022.154541

Suárez-Vidal, E., Sampedro, L., & Zas, R. (2017). Is the benefit of larger seed provisioning on seedling performance greater under abiotic stress? Environmental and Experimental Botany, 134, 45–53. 10.1016/j.envexpbot.2016.11.001

Sundqvist, M. K., Sanders, N. J., & Wardle, D. A. (2013). Community and Ecosystem Responses to Elevational Gradients: Processes, Mechanisms, and Insights for Global Change. Annual Review of Ecology, Evolution, and Systematics, 44(1), 261–280. 10.1146/annurev-ecolsys-110512-135750

Tasser, E., & Tappeiner, U. (2002). Impact of land use changes on mountain vegetation. Applied Vegetation Science, 5(2), 173–184. 10.1111/j.1654-109X.2002.tb00547.x

Van Kleunen, M., Weber, E., & Fischer, M. (2010). A meta-analysis of trait differences between invasive and non-invasive plant species. Ecology Letters, 13(2), 235–245. 10.1111/j.1461-0248.2009.01418.x

Violle, C., Garnier, E., Lecoeur, J., Roumet, C., Podeur, C., Blanchard, A., & Navas, M.-L. (2009). Competition, traits and resource depletion in plant communities. Oecologia, 160(4), 747–755. 10.1007/s00442-009-1333-x

Von Holle, B., Delcourt, H. R., & Simberloff, D. (2003). The importance of biological inertia in plant community resistance to invasion. Journal of Vegetation Science, 14(3), 425–432. 10.1111/j.1654-1103.2003.tb02168.x

Wagner, V., Chytrý, M., Jiménez-Alfaro, B., Pergl, J., Hennekens, S., Biurrun, I., Knollová, I., Berg, C., Vassilev, K., Rodwell, J. S., Škvorc, Ž., Jandt, U., Ewald, J., Jansen, F., Tsiripidis, I., Botta-Dukát, Z., Casella, L., Attorre, F., Rašomavičius, V., … Pyšek, P. (2017). Alien plant invasions in European woodlands. Diversity and Distributions, 23(9), 969–981. 10.1111/ddi.12592

Westoby, M., Falster, D. S., Moles, A. T., Vesk, P. A., & Wright, I. J. (2002). Plant ecological strategies: Some leading dimensions of variation between species. Annual Review of Ecology and Systematics, 33(1), 125–159.

WFO. (2024). World Flora Online. Published on: http://www.worldfloraonline.org. [Dataset].

World Meteorological Organization. (2017). Technical regulations basic documents No. 2 Vol. 1—General meteorological standards and recommended practices. Genève.

Wright, I. J., Reich, P. B., Westoby, M., Ackerly, D. D., Baruch, Z., Bongers, F., Cavender-Bares, J., Chapin, T., Cornelissen, J. H. C., Diemer, M., Flexas, J., Garnier, E., Groom, P. K., Gulias, J., Hikosaka, K., Lamont, B. B., Lee, T., Lee, W., Lusk, C., … Villar, R. (2004). The worldwide leaf economics spectrum. Nature, 428(6985), 821– 827. 10.1038/nature02403

Zuur, A. F., Ieno, E. N., Walker, N. J., Saveliev, A. A., & Smith, G. M. (2009). Mixed effects models and extensions in ecology with R (Vol. 574). Springer.

