## Supplementary for "Native community resistance modulates the spread of non-native species along Mediterranean mountain roads under global change"

**SUPPLEMENTARY MATERIAL**

The following Supplementary Material is available for this article:

**Table S1** Detailed description of the land cover types analysed with their relative attribution to the CORINE land cover classes.

**Table S2** Meta- and sub-models fitted in the SEM for testing the drivers of non-natives occurrence and non-natives abundance

**Table S3** List of non-native species recorded in the study area

**Table S4** Type III Analysis of Variance Table showing temporal changes in climate and land-use

**Table S5** Temporal changes in mean annual temperature (MAT, °C) and mean annual precipitation (MAP, mm)

**Table S6** Temporal changes in percentage landscape (PLand, %) across land-use cover types.

**Figure S1** Study area and sampling design

**Figure S2** SEM meta-model

**Table S1** Detailed description of the land cover types analysed with their relative attribution to the CORINE land cover classes.

| CORINE code | CORINE description | Detailed description | Class |
|---|---|---|---|
| 1. | Artificial surfaces | Artificial areas including: urban fabrics; industrial, commercial and transport units; mine, dump and construction sites; artificial non-agricultural vegetated areas. | ART |
| 1.2.2 | Road and rail networks | Include: road and railway transport network and associated lands. | ROA |
| 2. | Agricultural areas | Agricultural land, including: all types of arable land, permanent crops, pastures and heterogeneous agricultural areas. | AGR |
| 3.1 | Forests | Areas occupied by forests and woodlands with a vegetation pattern composed of native or exotic coniferous and/or broad-leaved trees | FOR |
| 3.1.2 | Afforestation | Vegetation formation composed principally of planted coniferous trees | AFF |
| 3.2.1 | Natural grasslands | Grasslands under no or moderate human influence. Low productivity grasslands. Often situated in areas of rough, uneven ground, steep slopes; frequently including rocky areas or patches of other (semi-)natural vegetation. | GRA |

**Table S2** Meta- and sub-models fitted in the SEM for testing the drivers of non-natives occurrence and non-natives abundance

| | **Models** |
|---|---|
| **SEM occurrence** | lmer(plant size ~ plot disturbance + current agricultural cover + current grassland cover + current artificial cover + current road cover +changes in agricultural cover + changes in grassland cover + changes in forest cover + current MAT + current MAP + changes in MAT + changes in MAP + (1\|mountain ID)) |
| | lmer(conservative-acquisitive strategy ~ plot disturbance + current agricultural cover + current grassland cover + current artificial cover + current road cover +changes in agricultural cover + changes in grassland cover + changes in forest cover + current MAT + current MAP + changes in MAT + changes in MAP + (1\|mountain ID)) |
| | lmer(functional diversity ~ plot disturbance + current agricultural cover + current grassland cover + current artificial cover + current road cover +changes in agricultural cover + changes in grassland cover + changes in forest cover + current MAT + current MAP + changes in MAT + changes in MAP + (1\|mountain ID)) |
| | glmer(non-natives presence/absence ~ plot disturbance + plant size + conservative-acquisitive strategy + functional diversity + current agricultural cover + current grassland cover + current artificial cover + current road cover +changes in agricultural cover + changes in grassland cover + changes in forest cover + current MAT + current MAP + changes in MAT + changes in MAP + (1\|mountain ID), family=binomial(link = "logit")) |
| **SEM abundance** | lmer(plant size ~ plot disturbance + current agricultural cover + current grassland cover + current artificial cover + current road cover +changes in agricultural cover + changes in grassland cover + changes in forest cover + current MAT + current MAP + changes in MAT + changes in MAP + (1\|mountain ID)) |
| | lmer(conservative-acquisitive strategy ~ plot disturbance + current agricultural cover + current grassland cover + current artificial cover + current road cover +changes in agricultural cover + changes in grassland cover + changes in forest cover + current MAT + current MAP + changes in MAT + changes in MAP + (1\|mountain ID)) |
| | lmer(functional diversity ~ plot disturbance + current agricultural cover + current grassland cover + current artificial cover + current road cover +changes in agricultural cover + changes in grassland cover + changes in forest cover + current MAT + current MAP + changes in MAT + changes in MAP + (1\|mountain ID)) |
| | lmer(non-natives cover (logit) ~ plot disturbance + plant size + conservative-acquisitive strategy + functional diversity + current agricultural cover + current grassland cover + current artificial cover + current road cover +changes in agricultural cover + changes in grassland cover + changes in forest cover + current MAT + current MAP + changes in MAT + changes in MAP + (1\|mountain ID)) |

**Table S3** List of non-native species recorded in the study area

| Species | Family | Growth form | Native area |
|---|---|---|---|
| *Aesculus hippocastanum* L. | Sapindaceae | Tree | West Asia |
| *Ailanthus altissima* (Mill.) Swingle | Simaroubaceae | Tree | East Asia |
| *Artemisia verlotiorum* Lamotte | Asteraceae | Perennial herb | East Asia |
| *Bromopsis inermis* (Leyss.) Holub subsp. *inermis* | Poaceae | Perennial grass | West Asia |
| *Cedrus deodara* (Roxb.) G.Don | Pinaceae | Tree | South Asia |
| *Ceratochloa cathartica* (Vahl) Herter | Poaceae | Perennial grass | South America |
| *Crepis sancta* (L.) Bornm. subsp. *nemausensis* (P.Fourn.) Babc. | Asteraceae | Annual herb | East Europe |
| *Erigeron annuus* (L.) Desf. | Asteraceae | Annual herb | North America |
| *Erigeron canadensis* L. | Asteraceae | Annual herb | North America |
| *Erigeron sumatrensis* Retz. | Asteraceae | Annual herb | Central America |
| *Euphorbia prostrata* Aiton | Euphorbiaceae | Annual herb | North America |
| *Hesperocyparis arizonica* (Greene) Bartel | Cupressaceae | Tree | North America |
| *Isatis tinctoria* L. | Brassicaceae | Annual herb | West Asia – North Africa |
| *Matricaria discoidea* DC. subsp. *discoidea* | Asteraceae | Annual herb | North America |
| *Robinia pseudoacacia* L. | Fabaceae | Tree | North America |
| *Senecio inaequidens* DC. | Asteraceae | Annual herb | South Africa |

**Table S4** Type III Analysis of Variance Table showing temporal changes in climate and land-use

| Response | Predictors | Sum of squares | DF | F | *P* |
|---|---|---|---|---|---|
| MAT | Time | 31.37 | 2, 118 | 704283 | **<0.001** |
| MAP | Time | 27313 | 2, 118 | 303.24 | **<0.001** |
| Land-use PLand | Time | 361 | 2, 531 | 0.37 | 0.691 |
| | Cover type | 310626 | 5, 563 | 127.25 | **<0.001** |
| | Time : Cover type | 20271 | 10, 527 | 4.15 | **<0.001** |

MAT, mean annual temperature; MAP, mean annual precipitation; PLand, percentage landscape

**Table S4** Temporal changes in mean annual temperature (MAT, °C) and mean annual precipitation (MAP, mm)

| | 1954 (T1) | | 1989 (T2) | | 2022 (T3) | |
|---|---|---|---|---|---|---|
| | mean | CI | mean | CI | mean | CI |
| MAT | $8.01^{a}$ | 0.54 | $7.98^{a}$ | 0.55 | $8.88^{b}$ | 0.55 |
| MAP | $769^{a}$ | 16 | $16^{a}$ | 122 | $16^{b}$ | 121 |

The table reports means and confidence intervals (CI) of MAT and MAP for the three climatic periods (1931-1960, 1961-1990, 1991-2020). Different superscripted letters indicate significant differences between periods (Tukey's post hoc test, *p-value* < 0.05).

**Table S6** Temporal changes in percentage landscape (PLand, %) across land-use cover types.

| | 1954 (T1) | | 1989 (T2) | | 2022 (T3) | |
|---|---|---|---|---|---|---|
| | mean | CI | mean | CI | mean | CI |
| Artificial areas | 5.86[a] | 14.22 | 8.01[a] | 8.93 | 8.61[a] | 8.47 |
| Roads | 3.76[a] | 6.53 | 3.85[a] | 5.99 | 3.50[a] | 6.04 |
| Agricultural areas | 52.32[a] | 8.77 | 31.56[b] | 9.90 | 18.08[b] | 11.30 |
| Afforestations | 12.88[a] | 13.59 | 24.55[a] | 9.89 | 27.1[a] | 10.13 |
| Forests | 53.35[a] | 7.85 | 62.12[ab] | 7.18 | 67.46[b] | 7.17 |
| Grasslands | 57.85[a] | 6.79 | 46.71[b] | 6.72 | 47.62[b] | 6.66 |

The table reports means and confidence intervals (CI) of the PLand of each land-use cover type for the three periods (1954, 1989, 2022). Different superscripted letters indicate significant differences between periods (Tukey's post hoc test, *p-value* < 0.05).

**Figure S1** Study area and sampling design

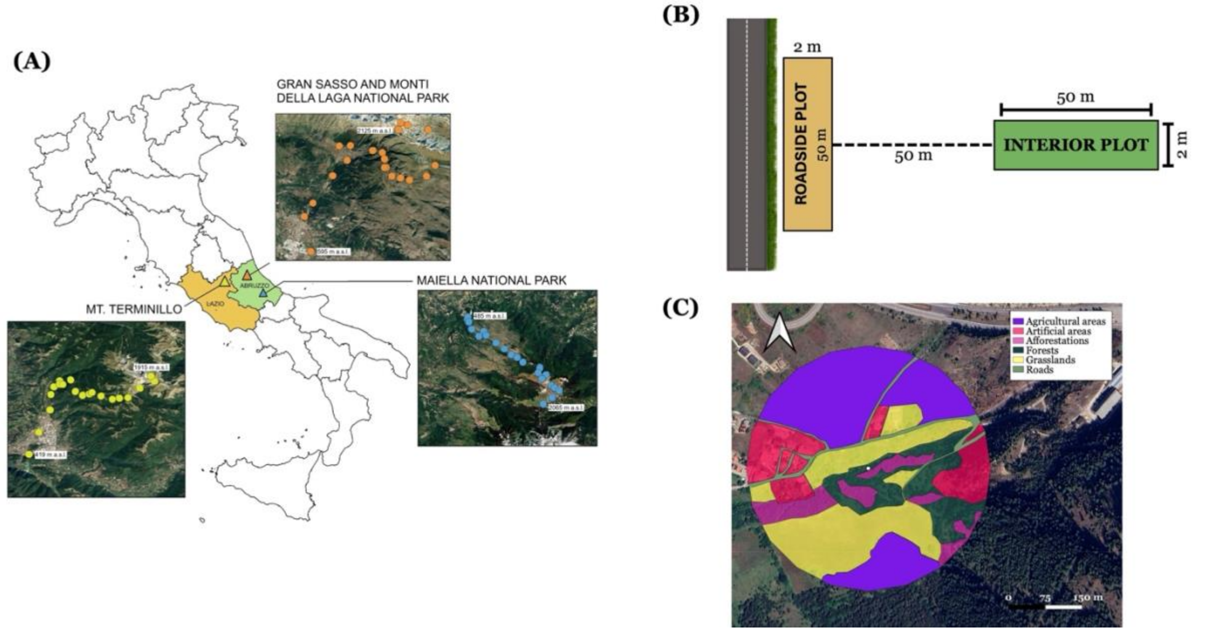


(**A**) Study area in the Central Apennines Mountain range (Italy) and the three mountain roads of Mount Terminillo, Mt. Maiella and the mountain chain of Gran Sasso. (**B**) Sampling design consisting of two 2 m × 50 m plots, one plot located parallel to the road (roadside plot) and a second plot perpendicular to the road and distant 50 m from the roadside (interior plot). (**C**) Example of the land-cover maps photo-interpreted within a circular buffer of 250 m of radius around the central point of each site. Figure adapted from (Santoianni et al., 2025)

**Figure S2** SEM meta-model

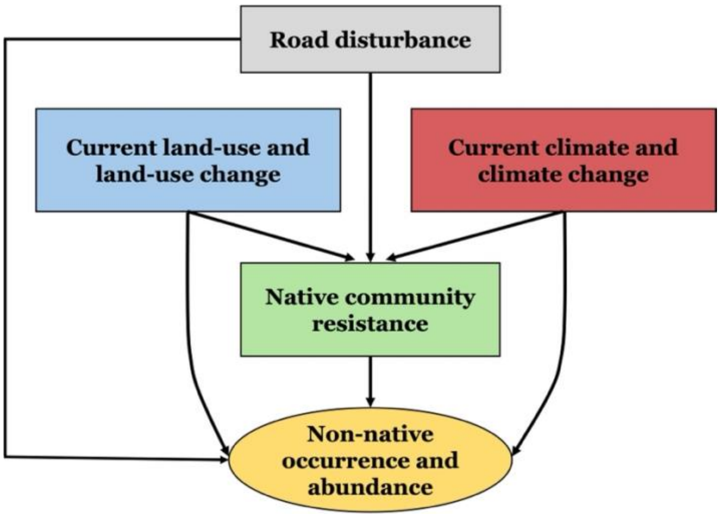
